# Heavy metals in moss guide environmental justice investigation: a case study using community science in Seattle, WA, USA

**DOI:** 10.1101/2022.04.20.488941

**Authors:** Sarah E. Jovan, Christopher Zuidema, Monika M. Derrien, Amanda L. Bidwell, Weston Brinkley, Robert J. Smith, Dale Blahna, Roseann Barnhill, Linn Gould, Alberto J. Rodríguez, Michael C. Amacher, Troy D. Abel, Paulina López

## Abstract

Heavy metals concentrations often vary at small spatial scales not captured by air monitoring networks, with implications for environmental justice in industrial-adjacent communities. Pollutants measured in moss tissues are commonly used as a screening tool to guide use of more expensive resources, like air monitors. Such studies, however, rarely address environmental justice issues or involve the residents and other decision-makers expected to utilize results. Here, we piloted a community science approach, engaging over 55 people from nine institutions, to map heavy metals using moss in two industrial-adjacent neighborhoods. This area, long known for disproportionately poor air quality, health outcomes, and racial inequities, has only one monitor for heavy metals. Thus, an initial understanding of spatial patterns is critical for gauging whether, where, and how to invest further resources towards investigating heavy metals. Local youth led sampling of the moss *Orthotrichum lyellii* from trees across a 250×250-m sampling grid (n = 79) and generated data comparable to expert-collected samples (n = 19). We mapped 21 chemical elements measured in moss, including 6 toxic ‘priority’ metals: arsenic, cadmium, chromium, cobalt, lead, and nickel. Compared to other urban *O. lyellii* studies, local moss had substantially higher priority metals, especially arsenic and chromium, encouraging community members to investigate further. Potential hotspots of priority metals varied somewhat but tended to peak near the central industrial core where many possible emissions sources, including legacy contamination, converge. Informed by these findings, community members successfully advocated regulators for a second study phase – a community-directed air monitoring campaign to evaluate residents’ exposure to heavy metals – as is needed to connect moss results back to the partnership’s core goal of understanding drivers of health disparities. This follow-up campaign will measure metals in the PM_10_ fraction owing to clues in the current study that airborne soil and dust may be locally important carriers of priority metals. Future work will address how our approach combining bioindicators and community science ultimately affects success addressing longstanding environmental justice concerns. For now, we illustrate the potential to co-create new knowledge, to help catalyze and strategize next steps, in a complex air quality investigation.

## 1. Introduction

The World Health Organization estimates that poor air quality causes 4.2 million premature deaths globally each year (Cohen et al. 2017). Understanding the spatial distributions of Hazardous Air Pollutants (HAPs; “air toxics”) is of international concern - even low levels of HAPs, including heavy metals, are associated with significant health risks such as various cancers, heart attack, stroke, asthma attacks, and neurological deficits (Jaishankar et al. 2014, Landrigan et al. 2018, USEPA 2014). According to the United States Government Accountability Office (2020), the ability to characterize HAPs at local scales is a critical need poorly supported by an aging air monitoring infrastructure. Declining funds and the high cost of regulatory-grade instruments makes for widely spaced monitoring networks, resulting in extensive areas of unknown exposure and environmental health risk (Marshall et al. 2008, Pakbin et al. 2010, Sarnat et al. 2010; Strum and Scheffe 2016).

To inexpensively fill these gaps for heavy metals, scientists commonly measure concentrations in moss or lichen tissue (hereafter, ‘bioindicators;’ e.g. Giordano et al. 2009, Massimi et al. 2019, Neitlich et al. 2017). Without roots or a protective outer cuticle, moss and lichens absorb water, nutrients, and co-occurring toxics from the atmosphere, making them a valuable first-pass screening tool in urban areas (e.g. Donovan et al. 2016, Messager et al. 2021). As with other low-cost sensors, bioindicator data help optimize the use of limited resources, like air monitors or even investigators’ time and attention (Donovan et al. 2016; Gatziolis et al. 2016). Air monitors are ultimately required to determine whether pollution levels pose human health risks or exceed regulatory thresholds, making their efficient use in air investigations essential.

Understanding fine-scale patterns may be particularly important in industrial-adjacent neighborhoods (United States Government Accountability Office 2020). Despite their demonstrated efficacy, however, bioindicators are rarely used in investigations of environmental justice (but see Contardo et al. 2018, Steiner et al. 2021), commonly defined as “the fair treatment and meaningful involvement of all people regardless of race, color, national origin, or income with respect to the development, implementation and enforcement of environmental laws, regulations and policies” (USEPA 2021). Furthermore, affected residents and other decision-makers are not typically involved in bioindicator research even though many collaborative environmental health studies report higher synergy, creativity, and efficiency, ultimately translating into greater research impacts (Cordner et al. 2019, English et al. 2018). By adopting a community science approach that substantively engages stakeholders, two distinct opportunities are created – for community engagement in the participatory process of environmental research itself, and in co-producing knowledge directly informing decisions that improve environmental conditions (Charles et al. 2020).

In this case study, community leaders convened a diverse partnership of public and non-profit organizations, led by residents and including local youth, to co-produce new knowledge addressing environmental justice concerns in the industrial-adjacent Georgetown and South Park neighborhoods in Seattle, WA, USA. Both are disproportionately burdened with poor health outcomes, air quality, and racial inequities, relative to other Seattle neighborhoods (Daniell et al. 2013, Gould and Cummings 2013, Min et al. 2019, Schulte et al. 2015, U.S. Department of Health and Human Services 2008, Washington State Department of Health 2021). Prior data suggest heavy metals may increase local cancer and non-cancer risk (Puget Sound Clean Air Agency (hereafter ‘PSCAA’) and Washington State Department of Ecology 2003, U.S. Department of Health and Human Services 2008, USEPA 2019) although only one monitor in the area measures heavy metals (Figure 1). Our study objectives included: 1) determining whether community and expert-collected moss samples indicate similar pollution patterns, 2) comparison with other urban moss datasets as context for local values, and 3) mapping and summarizing spatial distributions of heavy metals in moss. We discuss how results guide the course of investigation and directly informed three types of community action, including the successful initiation of a follow-up air monitoring campaign.

**Figure 1:**
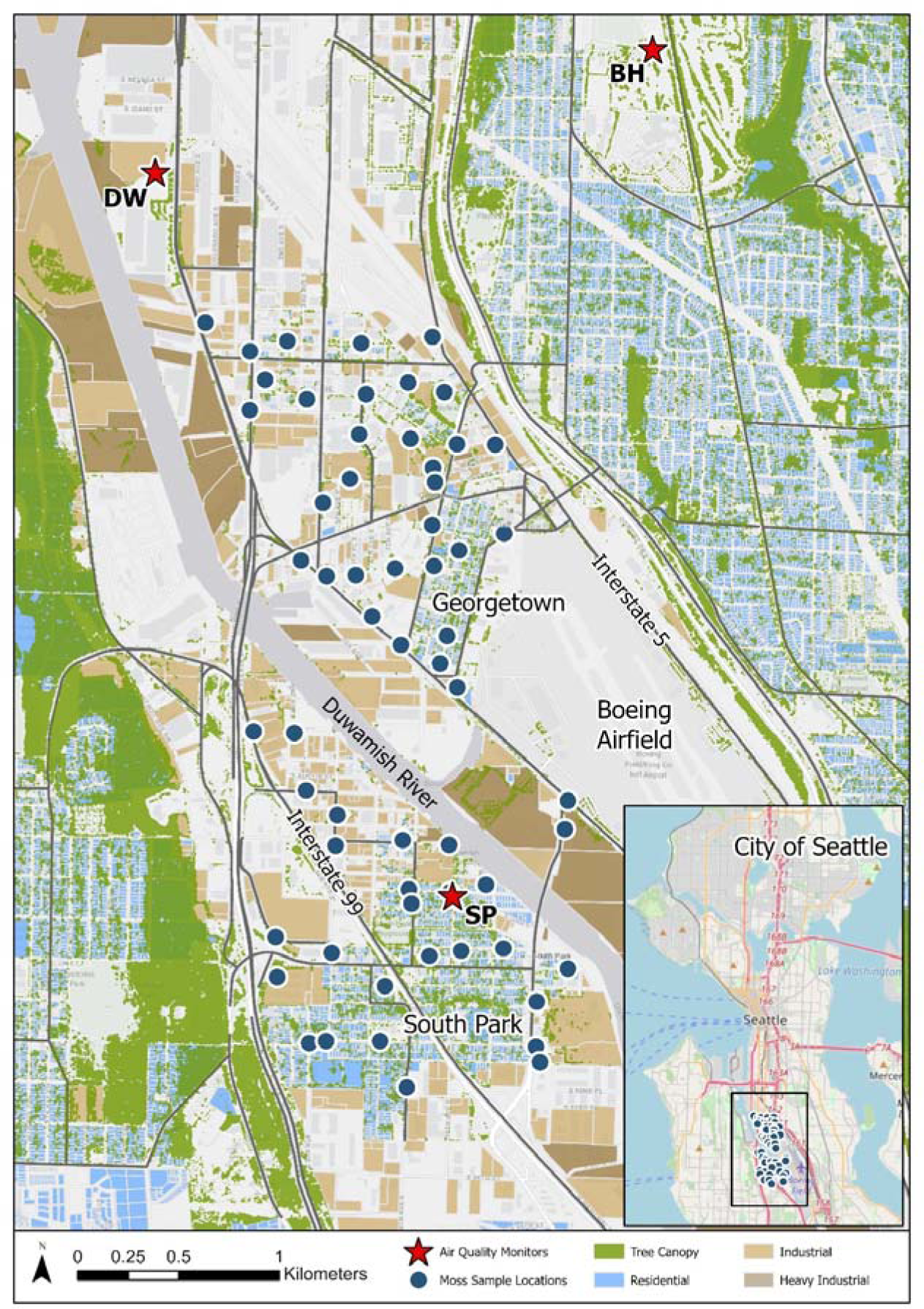
Map of the Duwamish Valley showing moss sampling locations and air quality monitors relative to major land use types (DW = Duwamish monitor, measures speciated PM_2.5_; BH = Beacon Hill, National Air Toxics Assessment site; includes speciated PM_2.5_ and PM_10_; SP = South Park monitor, total PM_2.5_ only).

## 2. Methods

### 2.1 Case Study Background

The South Park and Georgetown communities are located on the shores of the Duwamish River in the Duwamish Valley (DV) airshed (Figure 1). The Duwamish River has been Seattle’s main industrial corridor since the early 20^th^ century (Cummings 2020), which has created a legacy of potentially harmful pollution to air, soils and water. In 2001, a five-mile segment of the river, parts of which are adjacent to study area, was designated an active USEPA Superfund site due to contamination by polychlorinated biphenyls, carcinogenic polycyclic aromatic hydrocarbons, dioxins and furans, and arsenic in river sediments (USEPA 2013). In addition, over 150 Washington Model Toxics Control Act (MTCA; state “superfund”) contaminated sites have been named within or just adjacent to the study area, with clean-up completed at about 25% of them (Washington Department of Ecology 2021). The area also includes many unpaved roads, railway lines, waterway traffic, an airport (KCIA), highways with high levels of commuter and truck traffic, and several types of industrial facilities – some in place for over 100 years that manufacture or recycle materials such as glass, metal, and cement (Cummings 2020). These industrial and transportation areas are interspersed with single- and multi-family housing, home to approximately 5,600 Georgetown and South Park residents (City of Seattle 2018).

The long history of industrial pollution in Seattle is linked to poor health outcomes for residents (City of Seattle 2018, Gould and Cummings 2013, Min et al. 2019, Washington Department of Health 2021). Major health concerns include higher rates of asthma and diabetes, and lower life expectancies in the DV than city averages (City of Seattle 2018, Gould & Cummings 2013). Environmental factors map onto social vulnerabilities, resulting in cumulative health impacts that are higher than the rest of Seattle (Gould and Cummings 2013). In the 98108 ZIP code including Georgetown and South Park, 73.8% of residents are non-white (compared to 36.2% in Seattle city-wide), 34.7% are foreign-born (18.5% city-wide), 20.9% live below the poverty level (11.0% citywide), and just 32.2% of residents 25 years and older hold a bachelor’s degree or higher (64.0% city-wide; US Census 2019).

#### 2.1.1 Monitoring Sites and Standards

Heavy metals associated with particulate matter (PM) include fine inhalable particles with ≤ 2.5μm (i.e. PM_2.5_), inhalable particles with a diameter size ≤ 10 μm (PM_10_), and total suspended particulates (TSP). PM_10_ is considered hazardous to human health, with the finer fractions posing a higher risk due to deeper airway penetration (Brown et al. 2013). Both PM_2.5_ and PM_10_ are criteria pollutants, meaning their mass concentration in the atmosphere is regulated by federal ambient air standards. Similar standards do not exist for specific heavy metals except lead (Pb) in TSP.

There are currently 2 PM monitoring stations in the Duwamish Valley: the Duwamish (DW) site northwest of our moss sampling locations, and the South Park (SP) site at the north end of that neighborhood (Figure 1). Both DV stations measure the PM_2.5_ size fraction only, with “speciated” data (i.e., specifying concentrations of constituent heavy metals and other co-occurring toxics) available for just the DW site. The closest PM_10_ measurements come from Seattle’s single National Air Toxics Trends Site (NATTs) in the Beacon Hill (BH) neighborhood, about 1.5 km away above the DV river basin. This site measures many toxics, including heavy metals, in both PM_2.5_ and PM_10_, and is used as a reference site to broadly represent air quality across Seattle’s urban residential areas (PSCAA 2019). As larger particles (i.e. TSP) tend to be lower risk for health consequences, they are not criteria pollutants and are not often monitored outside of special studies.

### 2.2 Study Partners and Roles

Over 55 people from nine institutions, including community-based organizations, universities, and government agencies, played key roles in this project (See *Acknowledgements* section). Partnerships and collaborative process are detailed in Derrien et al. (2020). In brief, two organizations led youth engagement: the Duwamish Valley Youth Corps (DVYC), a program of the Duwamish River Community Coalition (DRCC) that engages high-school-aged youth in paid local environmental-justice oriented projects; and the Duwamish Infrastructure Restoration Training program (DIRT Corps), a green infrastructure workforce training program for adults. Study partners across institutions contributed to all facets of the study initiation, conception, sampling design, protocol development, youth engagement, data collection, and sample preparation (Derrien et al. 2020). Data analysis and interpretation was completed by agency and university partners.

### 2.3 Field Sampling and Preparation

Moss sampling protocols were adapted from prior studies using the widespread epiphytic (i.e. “tree-dwelling”) moss species *Orthotrichum lyellii* (Donovan et al. 2016, Gatziolis et al. 2016).

We collected the moss from a 250×250-m grid to ensure samples were well-distributed, although sample coverage along riverside industrial areas was sparser than intended due to lack of trees and target moss (Figure 1). Moss was collected at the suitable tree nearest the centroid of each grid cell. We used a “train-the-trainer” approach where scientists experienced in leading moss studies trained leaders of the DVYC and DIRT Corps, who then trained youth participants (Derrien et al. 2020). The DVYC led moss sampling excursions on four warm, dry days in 2019 (May 25^th^, June 1^st^, 4^th^, and 8^th^), working in five teams led by 3-5 youth, each accompanied by an adult study partner. Participants met the following week in a local high school science laboratory to harvest the upper 2/3^rds^ of living moss stems for heavy metals analysis (Gatziolis et al. 2016).

#### 2.3.1 Sampling QC/QA

To assess sampling precision, the youth-led teams immediately collected a replicate moss sample at 18 sites where ample moss was available. Their final analytical dataset had 79 samples from 61 grid cells. As reported initially in Derrien et al. (2020), measurements of heavy metals concentrations among same-day youth replicates were highly repeatable. To check sampling accuracy, experts re-sampled 19 grid cells although this occurred about 2 weeks later on June 13, 2019 due to scheduling difficulties. While Derrien et al. (2020) found sufficient statistical agreement between youth-expert samples to support confident use of the youth’s dataset in this study, they noted priority metals were somewhat lower in expert samples. Here, we investigated further by examining how well metals concentrations in the two datasets agreed spatially (Section 2.5.2) and by reviewing information on timing and other circumstances potentially affecting sample collection.

### 2.4 Laboratory Preparation and Analysis

All moss samples were sealed and mailed to the US Forest Service Grand Rapids, MN laboratory where they underwent the same treatment. Samples were prepared for heavy metals analysis by oven drying at 40 °C for 24 h and homogenizing by grinding to a fine powder (IKA tube-mill, 1 min grinding time for each sample at 15,000 rpm). A 0.500-g subsample of each moss sample was processed using a modified microwave-assisted digestion with 10 mL concentrated HNO_3_ + 2 mL 30% H_2_O_2_ + 2 mL concentrated HCl (CEM, 2019). An overnight pre-digestion of the samples with added reagents was done at room temperature. Following the microwave-assisted digestion cycle, digests were transferred by rinsing with deionized water to 50 mL volumetric flasks, diluted to volume with deionized water, and filtered through 0.45-um membrane filters into plastic storage bottles prior to analysis. Concentrations of 25 elements in total were measured by inductively coupled plasma optical emission spectrophotometry (Thermo 7000 series dual-view (axial and radial) ICP-OES).

#### 2.4.1 Lab QC/QA

Quality control steps included use of method blanks, instrument calibration standards, instrument performance check standards, and reference lichen samples. The measurement quality objectives (MQOs) are the same as the confidence and tolerance levels that accompany each standard reference material or check standard certification sheet. Quality control/quality assurance (QC/QA) for air-quality sampling in our study followed the EPA Quality Assurance Guidance document (USEPA 2016). Details of the laboratory QC/QA steps are provided in Appendix A.

### 2.5 Data Analysis

We used the youth-collected dataset to map 21 elements in moss tissues, leaving out 4 macronutrients of limited relevance (Table 1). Analyses focused on six heavy metals (hereafter, “priority metals”) commonly associated with negative ecological and human-health effects: arsenic, cadmium, chromium, cobalt, nickel, and lead. All are considered high priority for urban areas nationally (USEPA 2014, Agency for Toxic Substances and Disease Registry 2019) and all but cobalt locally (PSCAA 2019, PSCAA and University of Washington 2010, U.S. Department of Health and Human Services 2008, Wu et al. 2011).

**Table 1:**
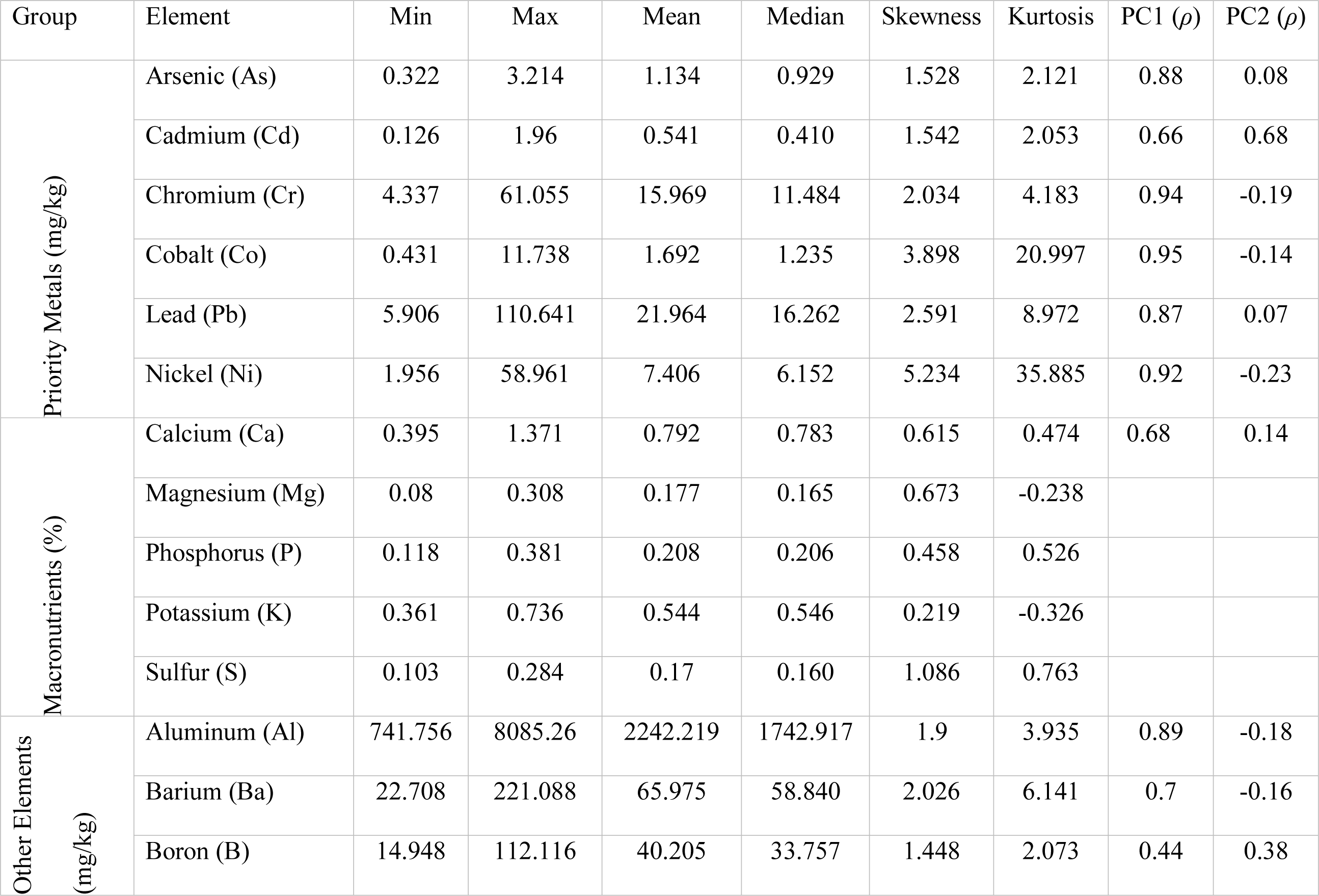
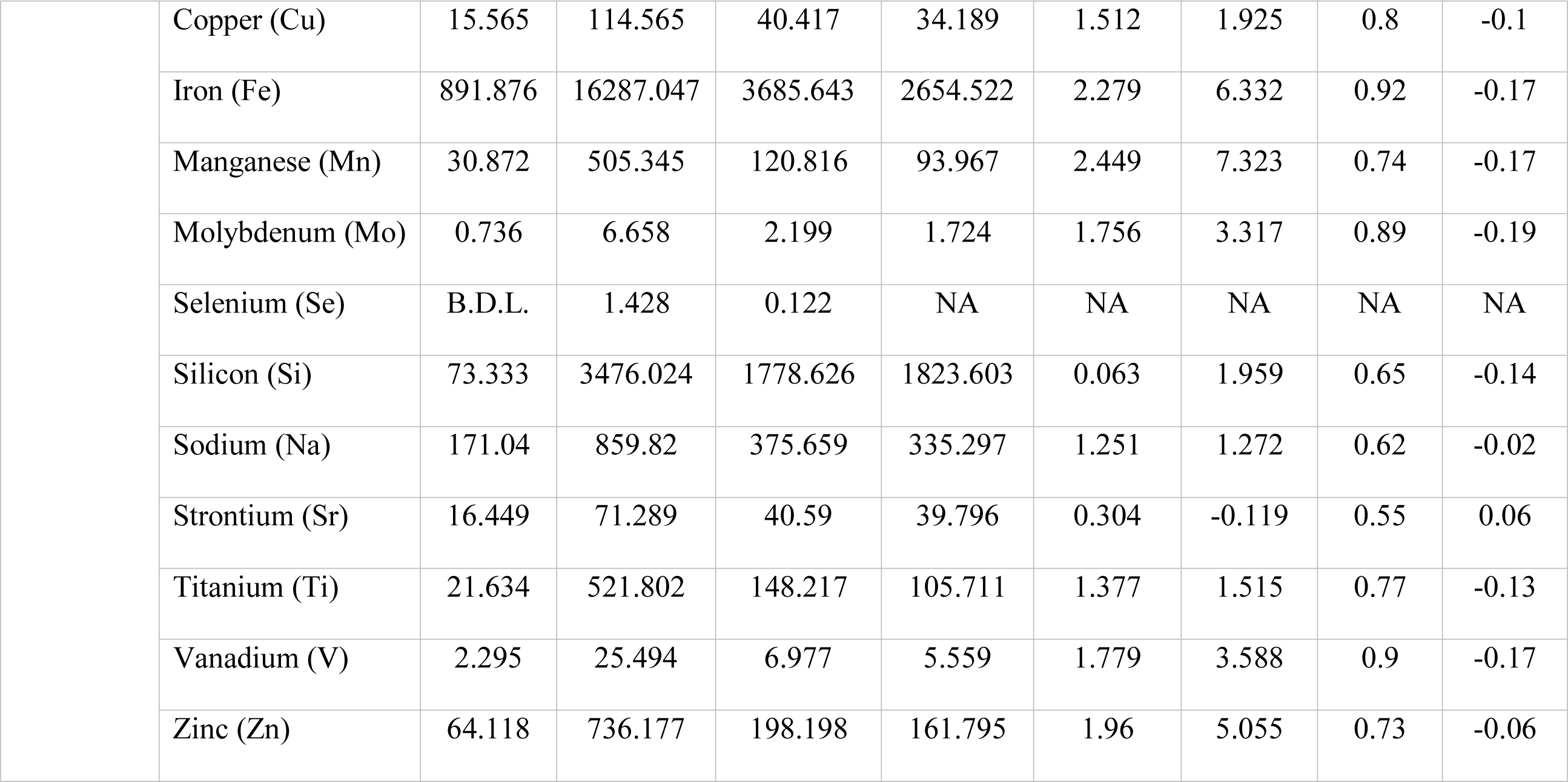
Summary of element concentrations in the youth’s moss collections (N = 79). Principal Component (PC) 1 and 2 are Spearman’s rank correlations between elements and axis scores from the first two axes found using Principal Components Analysis (PCA). Macronutrients of minor relevance were not included in the PCA. BDL = Below detection limit, NA = Not available.

| Group | Element | Min | Max | Mean | Median | Skewness | Kurtosis | PC1 ($\rho$) | PC2 ($\rho$) |
| --- | --- | --- | --- | --- | --- | --- | --- | --- | --- |
| Priority Metals (mg/kg) | Arsenic (As) | 0.322 | 3.214 | 1.134 | 0.929 | 1.528 | 2.121 | 0.88 | 0.08 |
|  | Cadmium (Cd) | 0.126 | 1.96 | 0.541 | 0.410 | 1.542 | 2.053 | 0.66 | 0.68 |
|  | Chromium (Cr) | 4.337 | 61.055 | 15.969 | 11.484 | 2.034 | 4.183 | 0.94 | -0.19 |
|  | Cobalt (Co) | 0.431 | 11.738 | 1.692 | 1.235 | 3.898 | 20.997 | 0.95 | -0.14 |
|  | Lead (Pb) | 5.906 | 110.641 | 21.964 | 16.262 | 2.591 | 8.972 | 0.87 | 0.07 |
|  | Nickel (Ni) | 1.956 | 58.961 | 7.406 | 6.152 | 5.234 | 35.885 | 0.92 | -0.23 |
| Macronutrients (%) | Calcium (Ca) | 0.395 | 1.371 | 0.792 | 0.783 | 0.615 | 0.474 | 0.68 | 0.14 |
|  | Magnesium (Mg) | 0.08 | 0.308 | 0.177 | 0.165 | 0.673 | -0.238 |  |  |
|  | Phosphorus (P) | 0.118 | 0.381 | 0.208 | 0.206 | 0.458 | 0.526 |  |  |
|  | Potassium (K) | 0.361 | 0.736 | 0.544 | 0.546 | 0.219 | -0.326 |  |  |
|  | Sulfur (S) | 0.103 | 0.284 | 0.17 | 0.160 | 1.086 | 0.763 |  |  |
| Other Elements (mg/kg) | Aluminum (Al) | 741.756 | 8085.26 | 2242.219 | 1742.917 | 1.9 | 3.935 | 0.89 | -0.18 |
|  | Barium (Ba) | 22.708 | 221.088 | 65.975 | 58.840 | 2.026 | 6.141 | 0.7 | -0.16 |
|  | Boron (B) | 14.948 | 112.116 | 40.205 | 33.757 | 1.448 | 2.073 | 0.44 | 0.38 |

|  |  |  |  |  |  |  |  |  |
| --- | --- | --- | --- | --- | --- | --- | --- | --- |
| Copper (Cu) | 15.565 | 114.565 | 40.417 | 34.189 | 1.512 | 1.925 | 0.8 | -0.1 |
| Iron (Fe) | 891.876 | 16287.047 | 3685.643 | 2654.522 | 2.279 | 6.332 | 0.92 | -0.17 |
| Manganese (Mn) | 30.872 | 505.345 | 120.816 | 93.967 | 2.449 | 7.323 | 0.74 | -0.17 |
| Molybdenum (Mo) | 0.736 | 6.658 | 2.199 | 1.724 | 1.756 | 3.317 | 0.89 | -0.19 |
| Selenium (Se) | B.D.L. | 1.428 | 0.122 | NA | NA | NA | NA | NA |
| Silicon (Si) | 73.333 | 3476.024 | 1778.626 | 1823.603 | 0.063 | 1.959 | 0.65 | -0.14 |
| Sodium (Na) | 171.04 | 859.82 | 375.659 | 335.297 | 1.251 | 1.272 | 0.62 | -0.02 |
| Strontium (Sr) | 16.449 | 71.289 | 40.59 | 39.796 | 0.304 | -0.119 | 0.55 | 0.06 |
| Titanium (Ti) | 21.634 | 521.802 | 148.217 | 105.711 | 1.377 | 1.515 | 0.77 | -0.13 |
| Vanadium (V) | 2.295 | 25.494 | 6.977 | 5.559 | 1.779 | 3.588 | 0.9 | -0.17 |
| Zinc (Zn) | 64.118 | 736.177 | 198.198 | 161.795 | 1.96 | 5.055 | 0.73 | -0.06 |

#### 2.5.1 Comparison to Reference Datasets

As heavy metals are one of several pollutant groups of concern in the DV, we conducted an initial screening step to gauge whether investigating them further was warranted. We compared priority metals to similar *O. lyellii* datasets from Seattle City Parks (n = 25; Bidwell 2018; Bidwell et al. 2019; Figure B.1) and residential areas throughout Portland, Oregon (n = 346; Donovan et al. 2016, Gatziolis et al. 2016). Sampled areas occur at similar elevations and human population densities in the “Dry Summer Subtropical Zone” of the Pacific Northwest (PNW), characterized by cool, wet winters and mild, relatively dry summers (Chen and Chen 2013). Datasets used similar field methods, digestion, and ICP spectroscopy techniques. One notable difference was sampling season; the DV was sampled in summer and reference datasets in winter. We used the 95^th^ percentiles for priority metals measured in Portland as points-of-comparison because these thresholds coincided with high outliers in that dataset, some of which exceeded local air concentration benchmarks when evaluated using air instruments.

#### 2.5.2 Spatial Distributions

We used Principal Components Analysis (PCA) based on the correlation structure among measured elements to identify and characterize major metals gradients in the youth’s dataset. We define ‘gradients’ as distinct distribution patterns involving correlated responses of multiple elements, represented by axes (i.e. principal components) in PCA analysis. First, we log_10_-transformed elements with highly skewed distributions (skewness > 0.5) to meet the normality and linearity assumptions of PCA. Second, we performed PCA on the scaled and centered correlation matrix of the six priority metals using function ’stats::prcomp’ in R version 4.0.2 (R Core Development Team 2020). Third, we applied graphical vector overlays and calculated Spearman’s rank correlation coefficients (ρ) to estimate how strongly all individual elements related to scores along each PCA axis (i.e. axis scores). Relationships between PCA axes and the other measured elements offer possible clues about the nature or origin of priority metals.

To evaluate spatial agreement between the youth- and expert-collected samples, we performed another set of PCA analyses separately for each dataset using only the subset of 17 sites having both youth and expert samples. To test agreement of youth and expert PCA scores, we compared axis scores using Procrustes analysis (Peres-Neto and Jackson 2001). We used the R function ’vegan::procrustes’ with symmetric solutions, calculated permutation *p*-values using 9999 permutations, and calculated Procrustean congruence (*R_P_*) as one minus the Procrustes sum-of-squared-errors (rather than the square-root of this quantity, as in software defaults) to better interpret it as a “coefficient of determination”-like statistic on a 0–1 scale. Perfect agreement among students’ and experts’ gradient scores would give a Procrustes fit (*R_P_*) approaching 1 and a permutation *p*-value << 0.01, while agreement no better than random would give *R_P_* near 0, and *p* >> 0.05.

Finally, we used the youth’s data to quantify the spatial distribution of pollution gradients by mapping scores from statistically significant PCA axes at moss sample locations, as well as by interpolating scores as a continuous surface between sites using spatial kriging. For kriging, we fit an empirical variogram model using the observed values to parameterize a final variogram model based on Gaussian covariance describing the spatial decay of similarity among sites (R functions ’gstat::variogram’ and ’gstat::vgm’). We assumed the Gaussian process was constant across the study area. From the variogram we applied ordinary kriging (’gstat::krigè) to predict interpolated values for unsampled locations between sites. Predicted values and their variances were used to construct 95% confidence intervals describing spatial uncertainty in the kriged PCA values. All spatial predictions were on a 10,000-cell grid in Albers equal-area projection covering the study area.

## 3. Results

Our final QC/QA check comparing PCA scores of youth vs expert-collected samples indicated highly significant agreement (*p*-value = 0.001) and sufficient spatial correlation (Procrustes fit *R_P_* = 0.42). Therefore, all remaining analyses are based solely on the youth’s data.

### 3.1 Comparison to Reference Datasets

Overall, priority metals in DV moss were significantly much greater than the Seattle City Parks and Portland residential datasets (t-tests; p <0.001). The most extreme cases were arsenic and chromium, for which nearly all DV concentrations exceeded Portland’s 95^th^ percentiles (Figure 2). Concentrations of cobalt, lead, nickel, and cadmium in the DV exceeded the Portland thresholds 59%, 55%, 43%, and 22% of the time, respectively. Moreover, medians and 25^th^ percentiles for DV samples were relatively elevated compared to reference datasets. The statistical sampling distributions of most elements in the DV were highly positively skewed, which typically indicates anthropogenic (vs natural geogenic) emissions sources (e.g. Solt et al. 2015; Figure 3, Figure B.2).

**Figure 2:**
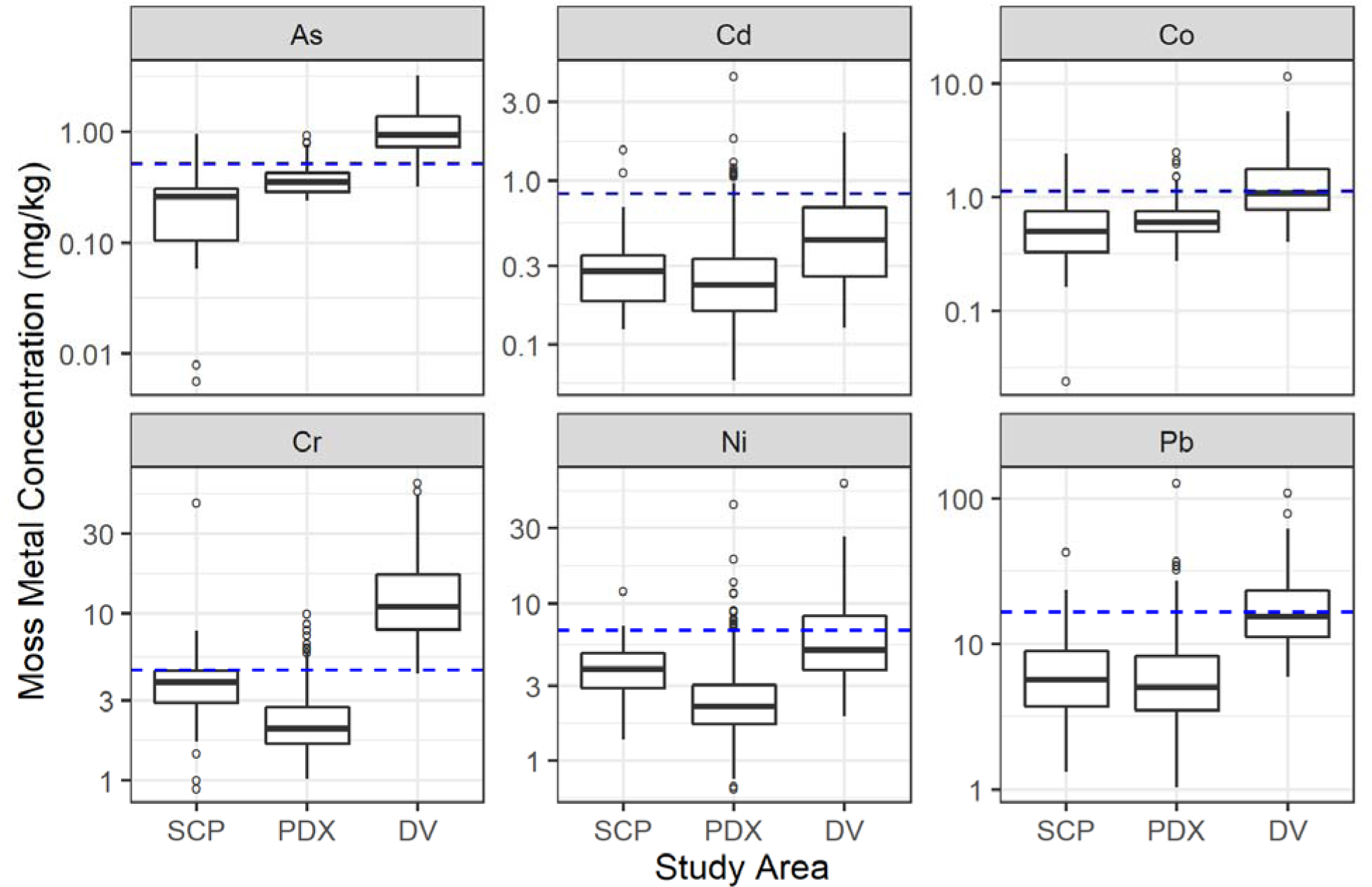
Boxplots comparing priority metals concentrations in this study (DV = Duwamish Valley) with Seattle-City Parks (SCP) and Portland-residential areas (PDX). Blue dashed lines indicate the Portland 95^th^ percentiles. Note that the y-axis is on the log scale.

**Figure 3:**
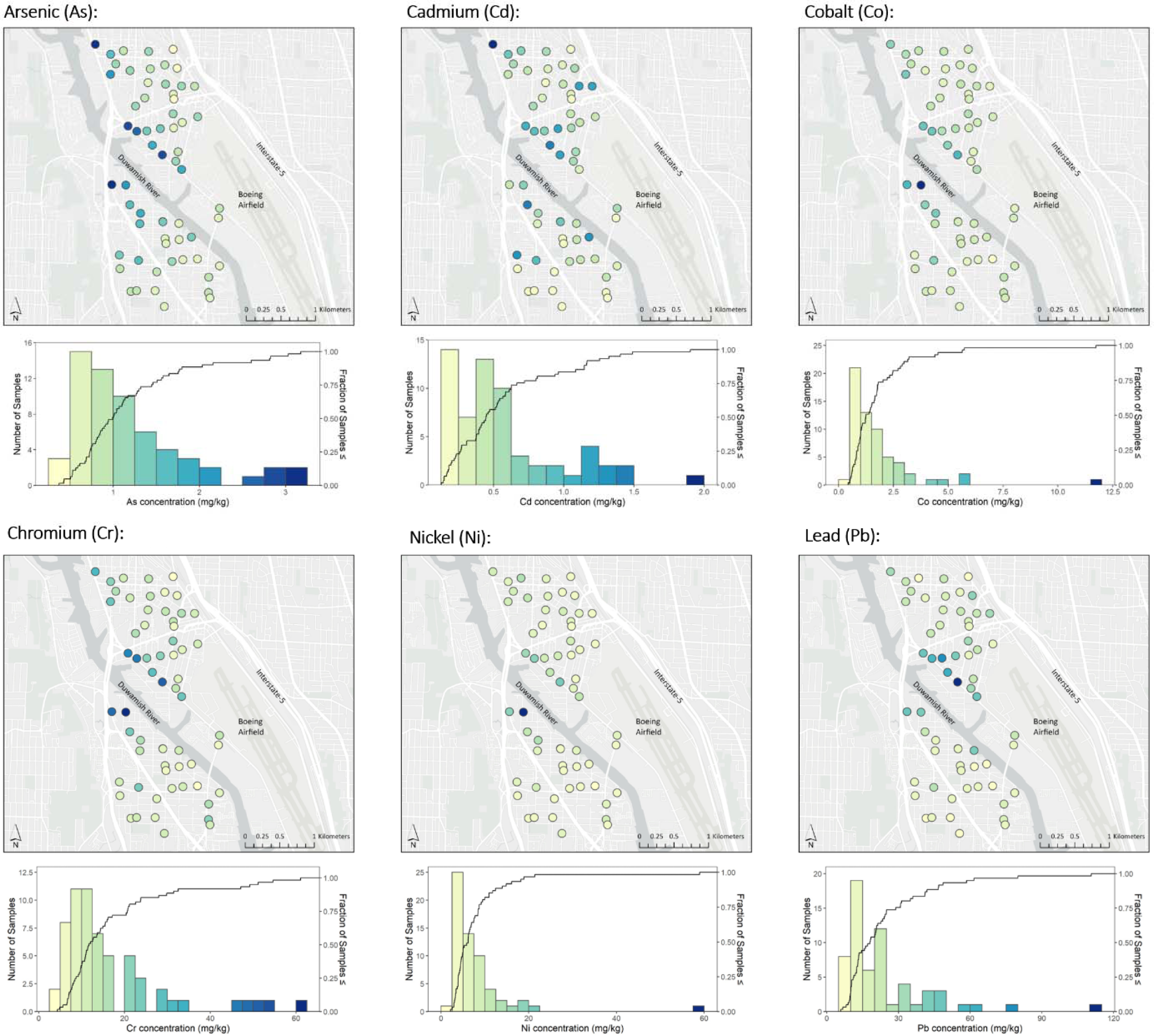
Dot maps and histograms showing concentrations of priority metals in DV moss. Black lines on the histograms are cumulative distribution curves.

### 3.2 Spatial Distributions

Concentrations of most elements peaked in the industrial core along the river’s banks west and northwest of Boeing Airfield and east of U.S. Highway 99, the main N-S highway crossing the river (Figure 3 & B.2). Priority metals were strongly correlated, such that sites with relatively high values for one tended to be high for the others (Figure B.3). The first PCA axis (‘PC1’, hereafter) supports this observation, explaining 76.5% of the variation in priority metals concentrations in the youth’s dataset (p = 0.001; Figure 4). We interpreted PC1 as the dominant gradient of elemental concentrations based on strong positive associations (Spearman’s correlations, r > 0.75) with nearly all measured elements (Table 1), indicating PC1 scores increased as elemental concentrations increased. Therefore, kriged PC1 scores shared common features with the maps for most individual metals (Figure 5). We also noted chemical elements indicative of soil and fugitive dust (i.e. aluminum, calcium, iron, silicon, strontium, titanium; Charlesworth et al. 2011, Kim and Hopke 2008, Watson and Chow 2000) correlated moderately to strongly with priority metals, besides cadmium, when considered both individually (mean Pearson correlation coefficient, r ≥ 0.65; Figure B.3) and collectively as PC1 (Spearman’s rank correlations, r = 0.55 to 0.92; Table 1).

**Figure 4.**
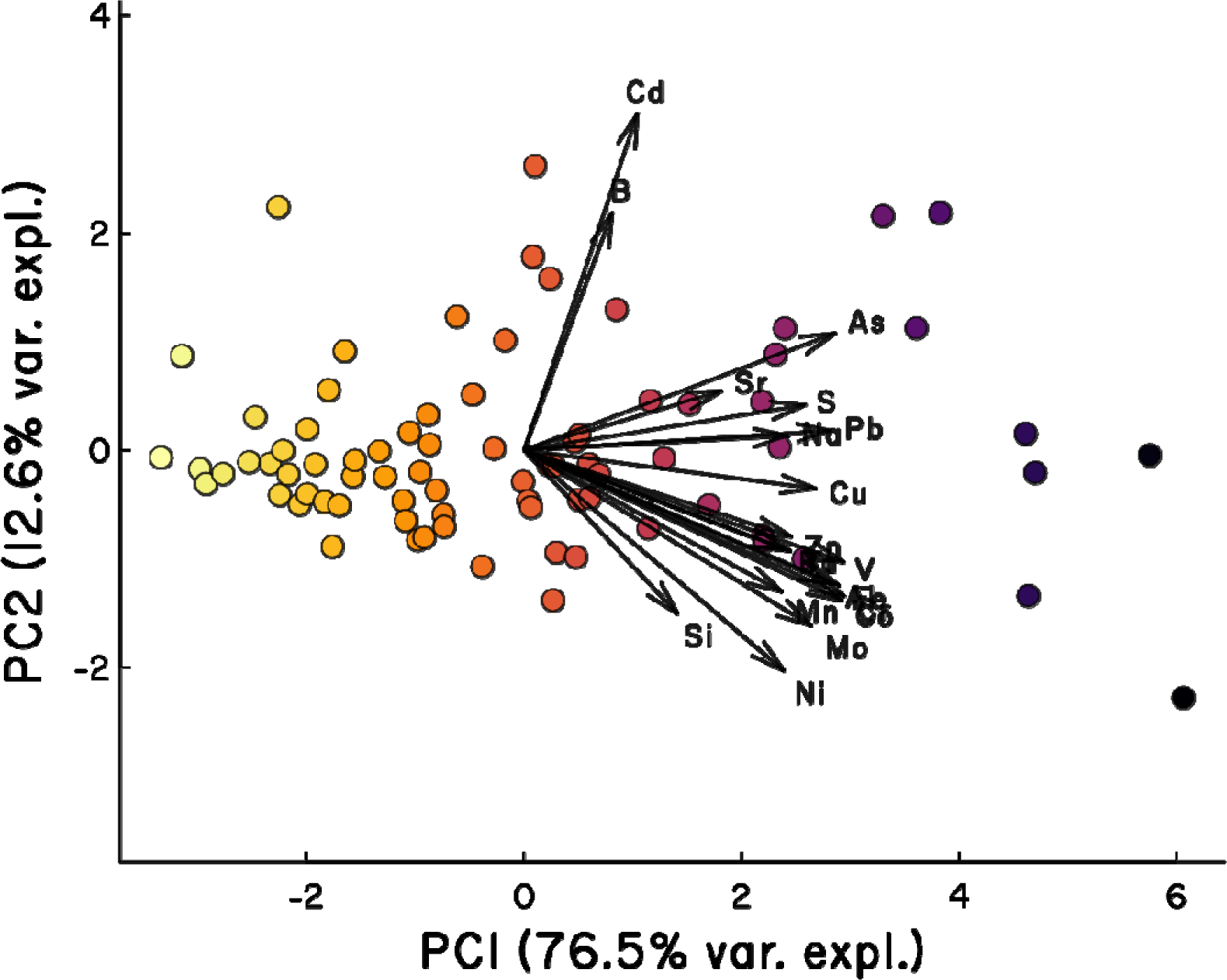
Biplot of scores from principal components analysis (PCA) of the six priority heavy metals (As, Cd, Cr, Co, Ni, Pb), with other measured elements shown as overlays. PCA axis 1 (‘PC1’) was the dominant axis of elemental concentration, with nearly all measured elements having strong positive correlations as indicated by direction and length of vector arrows. Each point represents one moss sample site, colored by relative value of PC1 axis scores (yellow = lowest to purple = highest). Concentrations were log_10_-transformed to linearize relationships and make highly skewed distributions more symmetrical.

**Figure 5:**
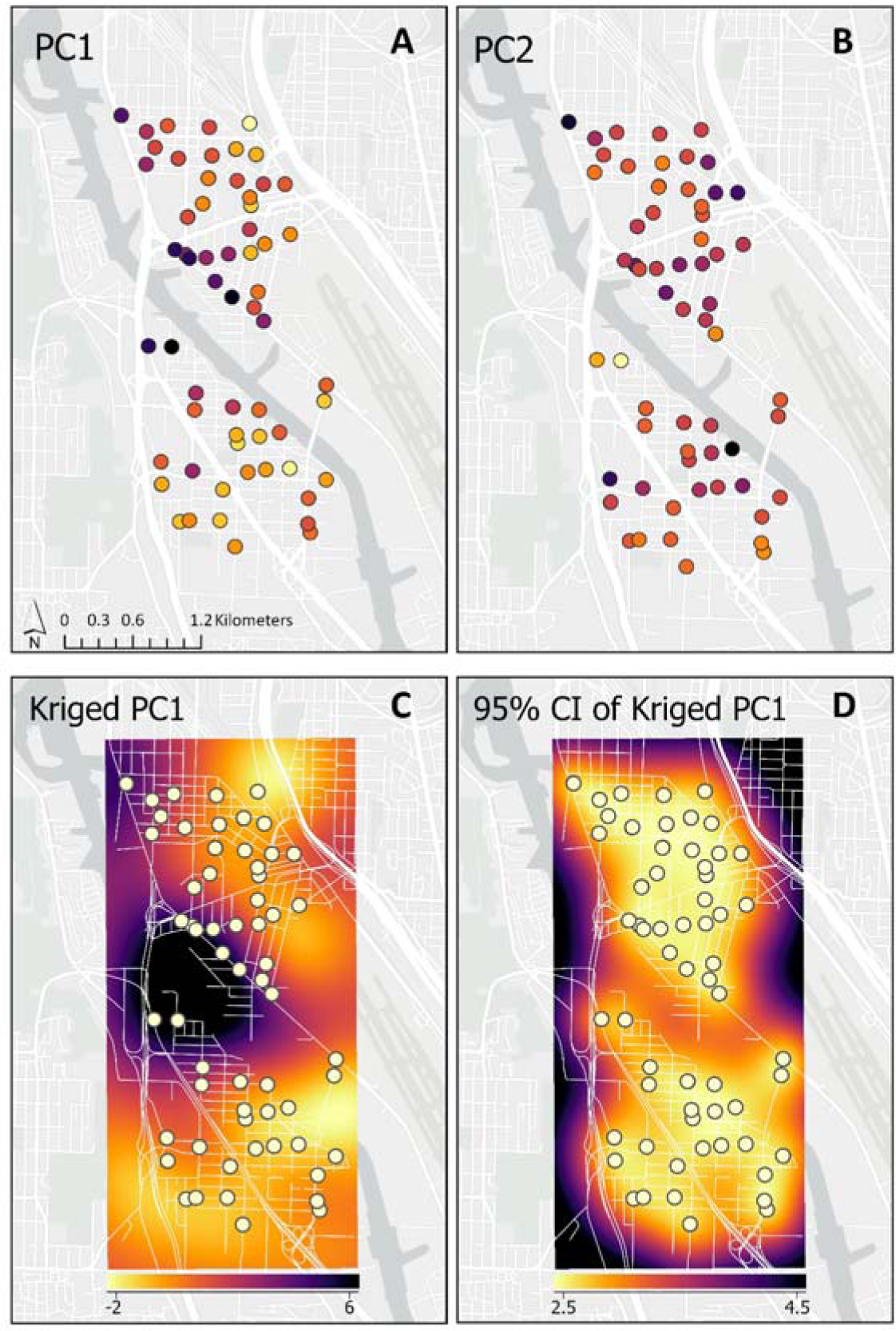
Maps of gradients in priority metals based on the similarity of elemental composition among moss sample sites. Each point, representing one sample site, is colored by its relative axis scores (yellow = lowest to purple = highest) along the two major gradients detected by PCA, PC1 (A) and PC2 (B). We used kriging to interpolate scores for the dominant gradient PC1 (C). Confidence intervals for kriged PC1 scores varied across the study area (D), showing the uncertainty of interpolated values was highest at the edges of the sampled area.

The second PCA axis (‘PC2’, hereafter) was non-significant, explaining only an additional 12.6% of the variation in priority metals (Figure 4). Therefore, we interpret it cautiously as a weak gradient with a unique elemental signature characterized by cadmium and boron concentrations (Spearman’s correlations r = 0.68 and 0.38, respectively). PC2 was also weakly positively associated with other elements like arsenic and strontium (Table 1).

## 4. Discussion

This was the first study of its kind in which residents, including local youth, collected and prepared moss samples for lab analysis with minimal oversight by scientific experts. Our results supported Derrien et al.’s (2020) conclusion that trained community and youth groups can collect scientifically viable moss tissue datasets. To further clarify Derrien et al.’s (2020) finding of somewhat lower priority metals concentrations in expert vs youth-collected samples, we confirmed these datasets captured similar spatial information when comparing relative rather than absolute concentrations. We suspect systematic differences were caused by a flash thunderstorm occurring after most youth but before most experts collected samples rather than a data quality issue. Conditions were mainly hot and dry during the three weeks of fieldwork, allowing PM to accumulate on bioindicator surfaces where driving rain can easily wash it off eburnis and Valiulis, 1999; Giordano et al. 2009).

### 4.1 Comparison to Reference Datasets

In our initial screening step comparing concentrations of priority metals in local and reference moss datasets, we interpreted our finding of much higher values in DV moss as strong justification for their further investigation. Our comparison is imperfect – for instance, datasets were collected in different seasons, which adds uncertainty to the comparison that is difficult to predict (e.g. Giordano et al. 2009, Saitanis et al. 2013). Nevertheless, we cannot easily assume weather or other unmeasured factors fully explain the large differences we observed (Figure 2). Furthermore, while spatially limited, prior studies using air monitors measured high priority metals concentrations at certain locales in the DV, including Georgetown (discussed in Section 4.3.1), which helped motivate the current study to characterize spatial patterns.

### 4.2 Spatial Distributions

The moss-based maps provide a first look at local-scale priority metals concentrations invaluable for guiding next steps of the investigation (Figures 3 & 5). The dense grid of moss samples shows high spatial variability that the single local air monitoring site (‘DW;’ Figure 1 & Figure B.1), or even a few hypothetical monitoring sites for that matter, couldn’t possibly describe. While our moss data (in mg/kg moss tissue) aren’t easily equated with air concentrations measured at DW, we noted the monitor did not detect arsenic at all (e.g. PSCAA 2018, 2019). This was surprising because several DV air investigations (e.g. King County 2015, PSCAA and Washington State Department of Ecology 2003, USEPA 2014, 2019) in addition to this study (Figure 2) suggest substantial arsenic concentrations in the DV. We present a testable hypothesis potentially explaining this discrepancy in Section 4.3.1, *Synthesis and Recommendations for Follow-up Air Monitoring*, and will continue focusing on arsenic in future work.

Overall, metals (as represented by PC1) tended to peak centrally where many extant emissions sources (industrial, highway, waterway, air travel) co-occur with legacy contamination of the soil accumulating over more than 100 years of industrial and transportation activity. Many unvegetated areas, including contaminated MTCA sites, exist near the hotspot along with unpaved roads where heavy truck traffic can track out dust and exacerbate re-entrainment into the air (Charlesworth et al. 2011, Roberts et. al 1975, Zhao et al. 2017). Accordingly, the close association of priority metals and elemental indicators of soil and fugitive dust (Section 3.2) suggests they may be part of the same particles or otherwise have common origins.

Efforts to identify specific sources affecting residential air quality lies beyond the scope of this study and will be complicated by the density of nearby emissions sources and potential for PM to continue mixing and affecting the air long after its initial emission (Johnston and Cushing 2020). How proximity to the central industrial area and other environmental factors correlate with metals levels in the neighborhoods’ residential zones (Figure 1) is the focus of a follow-up study already underway (Kondo et al. submitted). These results, along with findings from community actions focused on discerning the potential health risk of the metals (#3 in Section 4.3), will support decision-making on how to prioritize and approach unresolved questions about pollution sources.

The weak pattern indicated by PC2 seemed highly influenced by a single sample from the northwestern edge of the study area (Figure 5). This site had a unique elemental signature relatively high in cadmium, boron, arsenic and selenium (Table 1, Figure 3 & B.2 b,c,f,p) that was also detected in an expert re-sample collected a week later (data not shown). Follow-up work would be needed to understand the geographic scope and relevance of this finding. This unique signature could indicate one of several possibilities: a distinct emissions source just north of our sampling grid, an area-of-effect smaller than the resolution of our sampling grid, or simply idiosyncratic conditions at the particular tree where youth and adult study partners collected moss. Regardless, it’s clear PC2 does not describe a major pollution gradient in the current study area and is considered low priority for the community’s next steps investigating air concerns in Georgetown and South Park.

### 4.3 Community Actions

A core tenet of community science is that endeavors directly empower communities and support consequential collective actions (Charles et al. 2020). Partners’ ongoing attention to study design and outputs ensured that knowledge gaps being explored would result in actionable science addressing the specific needs of the Georgetown and South Park communities. Study processes and outputs resulted in three types of community action:

1. **Youth empowerment and training:** This work inspired youth actions in their communities, including mentoring other youth, leading independent data analysis, and presenting this study’s findings to community organizations including the mayor and city council. Youth engaged in and led many of the advocacy, partnership, and mitigation actions detailed below. Furthermore, the youth engaged in subsequent programming with project partners, including a second moss sampling campaign in summer 2021, expanding the sampling area and re-sampling areas of interest.
2. **Mitigation:** Our findings inform where to target near-term mitigation strategies to help improve conditions in the DV using green infrastructure (such as green walls and ongoing tree planting efforts) in partnership with local government and community organizations. Given racial inequities and health disparities in the DV, the timeliness of these near-term actions is especially important, and green infrastructure offers many immediate co-benefits. The implementation of mitigation strategies is also helping foster trust among the networks of project partners, critical for collaborative resource planning and management (Coleman & Stern 2018).
3. **Follow-up air monitoring:** Our findings equipped community partners with new knowledge about pollutant distributions at the neighborhood-scale. This helped community leaders advocate for a follow-up air monitoring campaign, in partnership with regulatory agencies (USEPA and PSCAA), to measure heavy metals in 2022. The campaign, sponsored by PSCAA, will deploy air monitoring instruments in locations selected by community partners based on local knowledge and the moss maps in this article (Figures 3, 5, B.2). Results will help determine whether metals detected in moss represent air concentrations with potential human health consequences. This critical question that will shape all subsequent steps in the partners’ heavy metals investigation.

#### 4.3.1 Synthesis and Recommendations for Follow-up Air Monitoring

After reviewing our results along with preexisting air modeling and monitoring data for the DV, we hypothesized that coarse particulates (PM_2.5-10_) may be important, overlooked carriers of arsenic and other heavy metals in the DV. First, most monitors measure PM_2.5_ only, including the local site ‘DW,’ which may explain why arsenic wasn’t detected there (Section 4.2). In contrast, moss tissues accumulate heavy metals within the broader PM_10_ size range (Adamo et al. 2008, Mariet et al. 2011, Massimi et al. 2019, Tretiach et al. 2011). Second, like the hotspots depicted in our maps, deposition of coarse particles is acute (i.e. concentrations vary widely across small areas) compared to the more gradual, regional-scale patterns of PM_2.5_ (e.g. Figure 5; Massimi et al. 2019, Pakbin et al. 2010, Zhang et al. 2014). Finally, airborne soil and dust (i.e. the main constituents of coarse particles; Watson and Chow 2000), clearly influenced the elemental signatures of moss samples (Section 4.2) and correlated closely with arsenic levels, a major known contaminant of local MCTA and DV Superfund sites.

To test our hypothesis, the follow-up air monitoring campaign will focus on measuring priority metals in PM_10_. Seattle’s only PM_10_ monitor is the BH NATTs site, a ‘reference’ monitor that by design would not capture local patterns (Goswami et al. 2002) in the DV where periodic stagnation events trap local pollution (Figure 1; Roberts et al. 1975, Su et al. 2008). Even so, it’s notable that in PM_10_ measured at BH, arsenic, cadmium, and chromium (as hexavalent chromium; Cr IV) rank among the top 12 HAPs with the highest potential cancer risk (PSCAA 2018, 2019). Our emphasis on needing to monitor local PM_10_ is underscored by prior studies showing significantly much higher arsenic, cadmium, and chromium in Georgetown versus BH in measurements of TSP and deposition flux (PSCAA and Washington State Department of Ecology 2003; King County 2015; Figure 1, B.1).

### 4.4 Study Limitations

We used a community science approach emphasizing the participation of youth and other community members, which required coordinating several groups with different time constraints. This led to three compromises in study design. First, field work spanned 3 weeks and a flash thunderstorm that appeared to affect the study objective of comparing youth vs expert re-samples (albeit not very importantly from a statistical perspective). Best practices for emphasizing spatial variability, however, are a brief (∼3 day) sampling window with consistent weather conditions. Second, we sampled in summer when lead partners were active, which wasn’t ideal for comparing with reference datasets collected in winter. Third, we sampled many street trees due to difficulty accessing interior trees on private lots, which is notable because particle deposition at roadside trees may be higher than trees short distances away (e.g. 100m; Pant and Harrison 2013). Nonetheless, we found strong and consistent geographic patterns of priority metals that helped partners prioritize and plan next steps with confidence – as was our goal for using moss as an inexpensive, first-pass screening tool.

## 5. Conclusions

As is common in urban neighborhoods, stakeholders in this study initially had few ‘on-the-ground’ measurements for evaluating the predominance and spatial distributions of heavy metals. Due to limited capacities for conventional air monitoring, supplemental datasets from low-cost sensors, such as bioindicators, may be greatly beneficial in complex airsheds like the DV where industrial and residential land commingle. As part of an on-going collaboration among many community, agency, and university partners, our study of bioindicators significantly catalyzed and advanced efforts to address longstanding environmental justice challenges, enabling new contributions to community-led problem solving related to air quality and health in the DV. Our main findings – that youth can collect high quality moss data; that priority metals in DV moss are relatively high vs reference datasets; that there are hotspots potentially warranting further investigation; that PM_10_ may be an important local carrier of metals we know little about – helped the partnership steer finite resources towards effective community actions. In addition to these main findings, our collaborative process built invaluable trust, social capital, and capacity among community and non-community research partners, serving as an important example of how community science partnerships and bioindicators can guide environmental health and justice work.

## Acknowledgements

Many partners were involved in this study. We gratefully acknowledge all 22 DVYC participants who led moss collection and preparation (Vanessa Arcos, Madison Brislin, Jose Camacho, Francisco Cano, Cristopher Castro, Alejandro Garcia, Adrian Gomez, Aalyiah Harren, Omar Lopez, Yaretzi Martinez, Nasheri Martinez, Deysi Olivera, Brendalyn Pastores, Joel Perez, Genevieve Perez, Mauricio Roman, Guadalupe Sanchez, Paola Silva, Cunningham Tach, Matthew Villalobos), with special thank you to Faith Villalobos and Leilani Guiterrez for helping present results to community groups and Seattle City Council members. Partners making significant contributions who aren’t co-authors include: Michelle C. Kondo (USFS N Research Station), Carmen Martinez (DRCC), Andrew Schiffer, Veronica Villarreal, and KC Steimer (DIRT Corps), Sandra Pinto Urrutia (City of Seattle Office of Sustainability & Environment), B.J. Cummings (University of Washington EDGE Center), Erik Saganić & Sarah Waldo (PSCAA), Karl Pepple (USEPA), Bruce McCune and Claudia Corvalan (Oregon State University), Abigail Kaminsky (USFS PNW Research Station), and Susan Will-Wolf (University of Wisconsin). A special thank you to John Larson, Randy Kolka, and Scott Kiel, who provided lab analysis of moss samples (USFS Grand Rapids Forestry Sciences Laboratory). Chris Zuidema was supported by the University of Washington’s Biostatistics, Epidemiology, and Bioinformatics Training in Environmental Health (BEBTEH), grant number T32ES015459, from the National Institute for Environmental Health Science (NIEHS). USDA Forest Service Voluntary Agreement 20-VI-11261985-0001 and Western Washington University professional leave fund supported Troy Abel. Interpretation and communication of results to the community was supported by the Community Engagement Core of the University of Washington Interdisciplinary Center for Exposures, Diseases, Genomics and Environment, NIEHS grant number P30ES007033. Project funding was also provided by the USDA Forest Service (State and Private Forestry Cooperative Agreement #18-CA-11062765-744, and Pacific Northwest Research Station Joint Venture Agreement #19-JV-11261985-072), the City of Seattle, Office of Sustainability & Environment, and the Urban Waters Federal Partnership, Green Duwamish designation through Street Sounds Ecology, LLC.

