## Supplemental Materials for "Heavy metals in moss guide environmental justice investigation: a case study using community science in Seattle, WA, USA"

Supplementary Materials

Appendix A

Quality Control and Quality Assurance

Quality control measures for the analysis of the moss samples included the use of reagent blanks (digestion reagents with no added elements), method blanks (reagent blanks carried through all digestion method steps), calibration standards diluted from commercially available multi-element concentrated ICP standards, and calibration verification and ICP performance check standards. ICP performance check standards were dilutions of commercially available concentrated method standards and included Alpha APS-1070 multi-element ICP standard, IQC-026 multi-element ICP standard (Agilent), VHG-IPC1Y multi-element ICP standard (LGC Standards), and QCI-710 water minerals standard (Agilent). Blanks and calibration checks were run every 10 sample digests.

To our knowledge, there is no reference moss sample with known analyte values. Instead, we used IAEA-336 reference lichen (*Evernia prunastri*) available from the International Atomic Energy Agency, and BCR-482 reference lichen (*Pseudevernia furfuracea*) available from the European Commission Joint Research Centre.

Results of the ICP performance check standards showed that the ICP was properly calibrated and capable of producing accurate values for the various analytes. Results for analysis of digests of the BCR-482 and IAEA-336 reference lichens are presented in appendix tables 1 and 2, respectively. Because all the Seattle moss samples were analyzed as a single set, the reported measured means ± standard errors in tables 1 and 2 are for 6 within-run subsamples. Since the Seattle study overlapped with a much larger Portland moss study, means ± standard errors for 13 separate Portland moss study analytical runs each consisting of several subsamples are reported in tables 1 and 2 for comparison with the Seattle data.

Measured values for the microwave-assisted HNO_3_ + H_2_O_2_ + HCl digestion method should be considered as total recoverable values rather than true totals. Our measured values for the major nutrient elements P, K, Mg, Ca, and S are slightly low for the BCR-482 lichen, whereas those for P and K in IAEA-336 lichen are closer to the confidence interval. IAEA-336 Mg and Ca results are informational values only, but our measured values are close to the listed values. For the trace nutrient elements Mo, Mn, Fe , Ni, Cu, Zn, and B, our measured values for Mo and Zn in BCR-482 lichen are low, are slightly low for Mn and B, and are within the uncertainties for Fe, Ni, and Cu. For IAEA-336 lichen, our measured values are slightly low for Zn and the informational value for Ni, but within the confidence intervals for Mn, Fe, and Cu. Reference values for Mo, and B are not given for the IAEA-336 lichen. For the soil mineral elements Na, Sr, Ba, Ti, Al, and Si in BCR-481 lichen, our measured values are low for Na, but only slightly low for Sr, Ba, Ti, and Al. No reference value was given for Si. For these same elements in IAEA-336 lichen, our measured values range from slightly low to within confidence intervals except for Ti where we had poor recovery.

Since a major goal of the Seattle moss study was to determine accumulation of environmentally important industrial-sourced trace metals (V, Cr, Co, Cd, Pb) and metalloids (As and Se), recovery of these elements in digests of the reference lichens was of paramount importance. For BCR-482 lichen, recoveries of all elements except Pb and Se were good. Our measured Pb values tended to be low while those for Se were slightly low. In contrast to the BCR-482 lichen data, all measured trace metals and metalloids in IAEA-336 were within the confidence intervals.

Table 1. ICP-AES quality control data for element analysis of BCR-482 *Pseudevernia* *furfuracea* reference sample (n = 13 separate analytical runs for Portland study, n = 1 analytical run for Seattle study, each run had multiple sub-samples). The P, K, Mg, Ca, S, Mo, Mn, Fe, B, Na, Sr, Ba, Ti, V, Co, and Se concentration values are not certified reference values and are for informational purposes only.

| Element  (unit) | BCR value | Uncertainty | Portland study  measured range | | | | Portland study  measured  mean ± std err | | | Seattle study  measured  mean ± std err | | | | Detection  limit (mg/kg)^[1]^ | |
| --- | --- | --- | --- | --- | --- | --- | --- | --- | --- | --- | --- | --- | --- | --- | --- |
| P (%) | 0.0690 | 0.0010 | 0.054 | - | 0.060 | 0.057 | | ± | 0.001 | | 0.059 | ± | 0.000 | | 0.155 |
| K (%) | 0.3900 | 0.0224 | 0.303 | - | 0.349 | 0.342 | | ± | 0.004 | | 0.351 | ± | 0.003 | | 0.060 |
| Mg (%) | 0.0578 | 0.0004 | 0.043 | - | 0.050 | 0.047 | | ± | 0.001 | | 0.048 | ± | 0.000 | | 0.001 |
| Ca (%) | 0.2624 | 0.0180 | 0.195 | - | 0.227 | 0.213 | | ± | 0.003 | | 0.206 | ± | 0.001 | | 0.000 |
| S (%) | 0.2166 | 0.0292 | 0.126 | - | 0.172 | 0.145 | | ± | 0.004 | | 0.163 | ± | 0.001 | | 0.105 |
| Mo (mg/kg) | 0.85 | 0.01 | 0.316 | - | 0.682 | 0.413 | | ± | 0.027 | | 0.393 | ± | 0.011 | | 0.038 |
| Mn (mg/kg) | 33.0 | 0.5 | 25.1 | - | 30.9 | 27.1 | | ± | 0.4 | | 24.9 | ± | 0.1 | | 0.007 |
| Fe (mg/kg) | 804 | 160 | 641 | - | 823 | 747 | | ± | 17 | | 741 | ± | 8 | | 0.025 |
| Ni (mg/kg) | 2.47 | 0.07 | 1.63 | - | 4.32 | 2.25 | | ± | 0.22 | | 1.89 | ± | 0.05 | | 0.036 |
| Cu (mg/kg) | 7.03 | 0.19 | 5.73 | - | 7.42 | 6.42 | | ± | 0.14 | | 6.08 | ± | 0.05 | | 0.039 |
| Zn (mg/kg) | 100.6 | 2.2 | 69.69 | - | 81.10 | 76.67 | | ± | 0.95 | | 78.28 | ± | 0.53 | | 0.019 |
| B (mg/kg) | 4.3 | 0.1 | 1.57 | - | 7.98 | 2.70 | | ± | 0.55 | | 2.60 | ± | 0.21 | | 0.069 |
| Na (mg/kg) | 119 | 2 | 45 | - | 61 | 54 | | ± | 1 | | 54 | ± | 1 | | 0.037 |
| Sr (mg/kg) | 10.35 | 0.24 | 8.0 | - | 9.3 | 8.8 | | ± | 0.1 | | 8.9 | ± | 0.0 | | 0.001 |
| Ba (mg/kg) | 14.9 | 2.4 | 9.5 | - | 13.4 | 10.9 | | ± | 0.3 | | 10.1 | ± | 0.2 | | 0.003 |
| Ti (mg/kg) | 34.2 | 1.1 | 17.2 | - | 46.9 | 30.6 | | ± | 1.9 | | 22.2 | ± | 1.9 | | 0.030 |
| Al (mg/kg) | 1103 | 24 | 657 | - | 925 | 806 | | ± | 21 | | 746 | ± | 38 | | 0.012 |
| Si (mg/kg) |  |  | 734 | - | 1313 | 1027 | | ± | 47 | | 1104 | ± | 62 | | 0.109 |
| V (mg/kg) | 3.74 | 0.61 | 3.03 | - | 4.17 | 3.51 | | ± | 0.08 | | 3.18 | ± | 0.05 | | 0.023 |
| Cr (mg/kg) | 4.12 | 0.15 | 3.04 | - | 6.85 | 3.75 | | ± | 0.29 | | 3.28 | ± | 0.14 | | 0.021 |
| Co (mg/kg) | 0.32 | 0.03 | 0.262 | - | 0.385 | 0.317 | | ± | 0.010 | | 0.281 | ± | 0.007 | | 0.051 |
| Cd (mg/kg) | 0.56 | 0.02 | 0.357 | - | 0.452 | 0.394 | | ± | 0.007 | | 0.440 | ± | 0.004 | | 0.007 |
| Pb (mg/kg) | 40.9 | 1.4 | 29.714 | - | 34.009 | 32.438 | | ± | 0.334 | | 32.423 | ± | 0.282 | | 0.106 |
| As (mg/kg) | 0.85 | 0.07 | 0.654 | - | 1.048 | 0.845 | | ± | 0.034 | | 0.979 | ± | 0.057 | | 0.143 |
| Se (mg/kg) | 0.6 | 0.2 | -0.057 | - | 0.746 | 0.350 | | ± | 0.063 | | 0.226 | ± | 0.056 | | 0.305 |

Table 2. ICP-AES quality control data for element analysis of IAEA-336 *Evernia* *prunastri* reference sample (n = 13 separate analytical runs for Portland study, n = 1 analytical run for Seattle study, each run had multiple sub-samples). The P, Mg, Ca, Ni, Al, Cr, Cd, and Pb concentration values supplied with the IAEA-336 sample are not recommended reference values and are for informational purposes only (Mg, Ca, and Ni are listed as uncertain).

| Element  (unit) | IAEA value  (95% CI) | Portland study  measured range | | | | Portland study  measured  mean ± std err | | | | Seattle study  measured  mean ± std err | | | Detection  Limit  (mg/kg)^[2]^ |
| --- | --- | --- | --- | --- | --- | --- | --- | --- | --- | --- | --- | --- | --- |
| P (%) | 0.061 (0.049-0.073) | 0.045 | - | 0.048 | 0.046 | | ± | 0.000 | 0.049 | | ± | 0.000 | 0.155 |
| K (%) | 0.184 (0.164-0.204) | 0.143 | - | 0.168 | 0.161 | | ± | 0.002 | 0.172 | | ± | 0.002 | 0.060 |
| Mg (%) | 0.058 | 0.048 | - | 0.055 | 0.053 | | ± | 0.001 | 0.055 | | ± | 0.000 | 0.001 |
| Ca (%) | 0.282 | 0.211 | - | 0.248 | 0.230 | | ± | 0.004 | 0.231 | | ± | 0.003 | 0.000 |
| S (%) | none listed | 0.047 | - | 0.081 | 0.055 | | ± | 0.003 | 0.063 | | ± | 0.000 | 0.105 |
| Mo (mg/kg) | none listed | -0.012 | - | 0.234 | 0.071 | | ± | 0.017 | 0.078 | | ± | 0.008 | 0.038 |
| Mn (mg/kg) | 63 (46-70) | 55.8 | - | 64.8 | 59.0 | | ± | 0.8 | 56.1 | | ± | 0.5 | 0.007 |
| Fe (mg/kg) | 430 (380-480) | 346 | - | 437 | 395 | | ± | 8 | 392 | | ± | 6 | 0.025 |
| Ni (mg/kg) | 1.65 | 0.46 | - | 1.09 | 0.79 | | ± | 0.05 | 0.80 | | ± | 0.05 | 0.036 |
| Cu (mg/kg) | 3.6 (3.1-4.1) | 2.53 | - | 4.47 | 3.15 | | ± | 0.14 | 3.09 | | ± | 0.06 | 0.039 |
| Zn (mg/kg) | 30.4 (27.0-33.8) | 22.86 | - | 28.18 | 24.73 | | ± | 0.43 | 25.63 | | ± | 0.28 | 0.019 |
| B (mg/kg) | none listed | 1.07 | - | 3.78 | 1.87 | | ± | 0.20 | 1.45 | | ± | 0.03 | 0.069 |
| Na (mg/kg) | 320 (280-360) | 212 | - | 294 | 265 | | ± | 6 | 280 | | ± | 4 | 0.037 |
| Sr (mg/kg) | 9.3 (8.2-10.4) | 6.9 | - | 8.3 | 7.8 | | ± | 0.1 | 8.2 | | ± | 0.0 | 0.001 |
| Ba (mg/kg) | 6.4 (5.3-7.5) | 4.5 | - | 5.8 | 5.1 | | ± | 0.1 | 4.9 | | ± | 0.2 | 0.003 |
| Ti (mg/kg) | 50 | 11.6 | - | 22.9 | 15.7 | | ± | 0.8 | 11.4 | | ± | 1.5 | 0.030 |
| Al (mg/kg) | 680 (570-790) | 461 | - | 588 | 528 | | ± | 11 | 506 | | ± | 34 | 0.012 |
| Si (mg/kg) | 1181 | 490 | - | 999 | 757 | | ± | 39 | 871 | | ± | 81 | 0.109 |
| V (mg/kg) | 1.47 (1.25-1.69) | 1.12 | - | 1.53 | 1.26 | | ± | 0.03 | 1.13 | | ± | 0.04 | 0.023 |
| Cr (mg/kg) | 1.06 (0.89-1.23) | 0.97 | - | 1.76 | 1.19 | | ± | 0.06 | 1.07 | | ± | 0.03 | 0.021 |
| Co (mg/kg) | 0.29 (0.24-0.34) | 0.222 | - | 0.598 | 0.303 | | ± | 0.032 | 0.254 | | ± | 0.002 | 0.051 |
| Cd (mg/kg) | 0.117 (0.100-0.134) | 0.076 | - | 0.125 | 0.096 | | ± | 0.004 | 0.103 | | ± | 0.004 | 0.007 |
| Pb (mg/kg) | 4.9 (4.3-5.5) | 4.379 | - | 6.515 | 4.887 | | ± | 0.182 | 4.661 | | ± | 0.025 | 0.106 |
| As (mg/kg) | 0.63 (0.55-0.71) | 0.597 | - | 0.932 | 0.748 | | ± | 0.032 | 0.724 | | ± | 0.072 | 0.143 |
| Se (mg/kg) | 0.22 (0.18-0.26) | -0.040 | - | 0.813 | 0.333 | | ± | 0.063 | 0.227 | | ± | 0.028 | 0.305 |

Appendix B

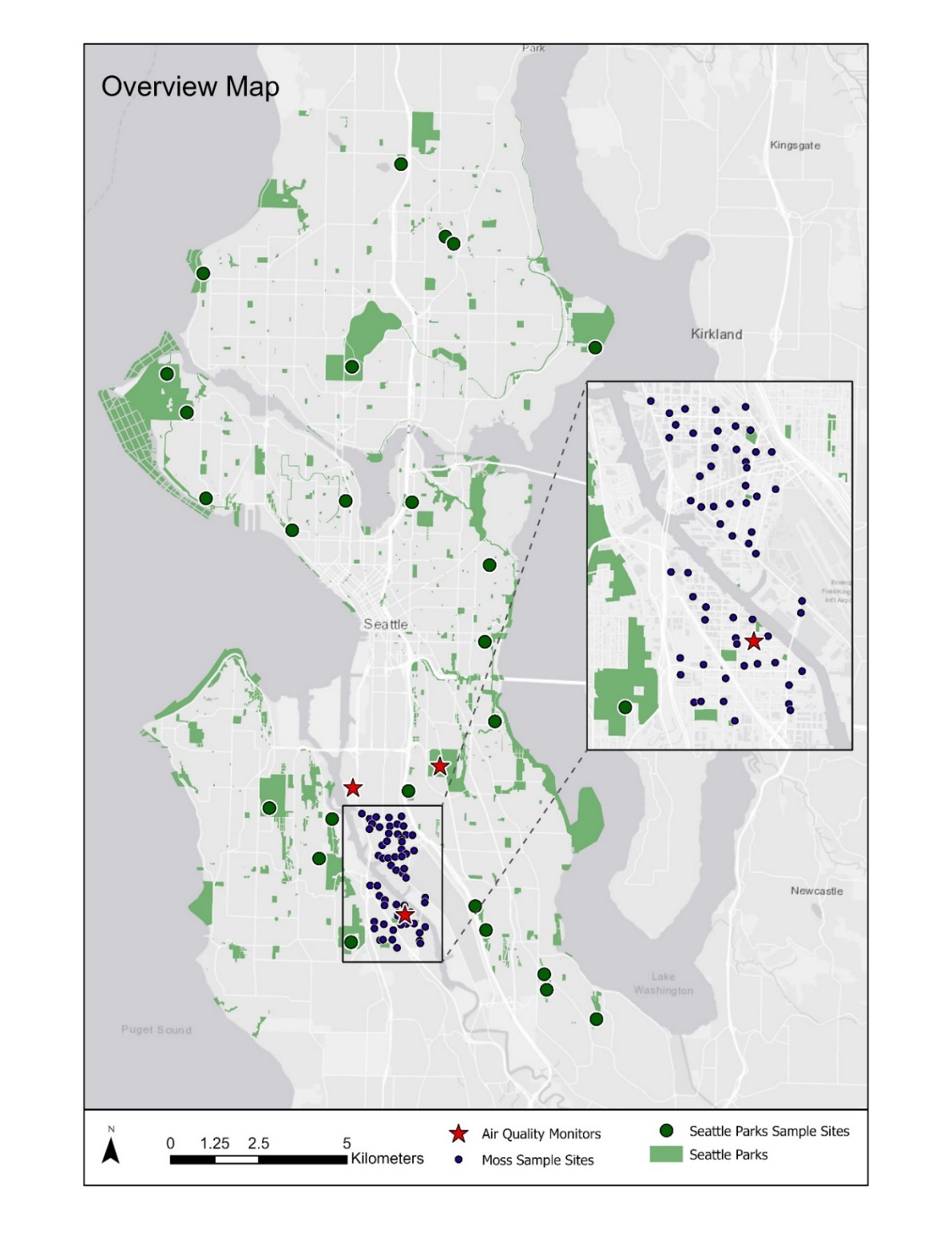

Figure B.1: Map showing moss sample sites in the Duwamish Valley and Seattle City Parks datasets.

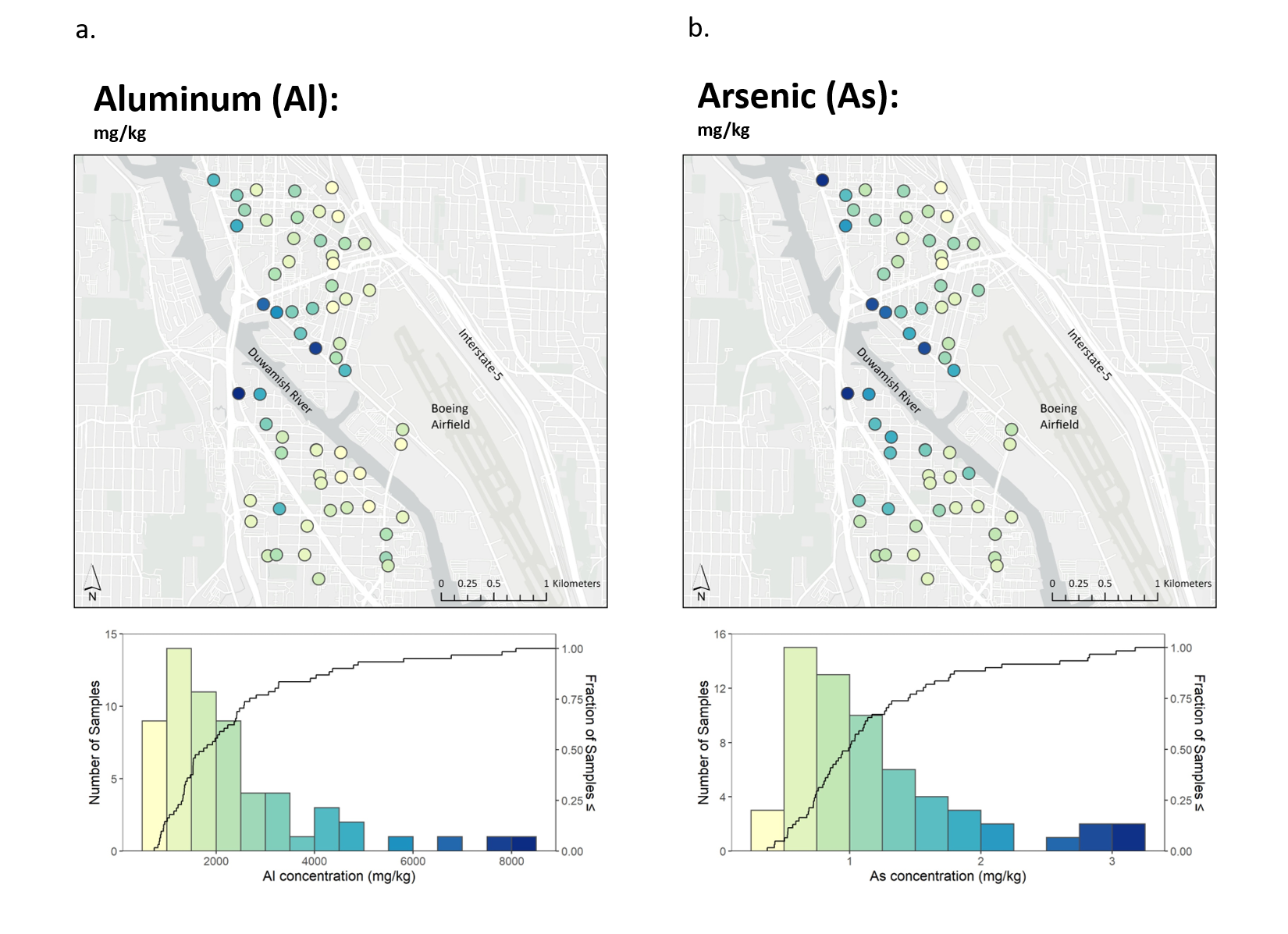

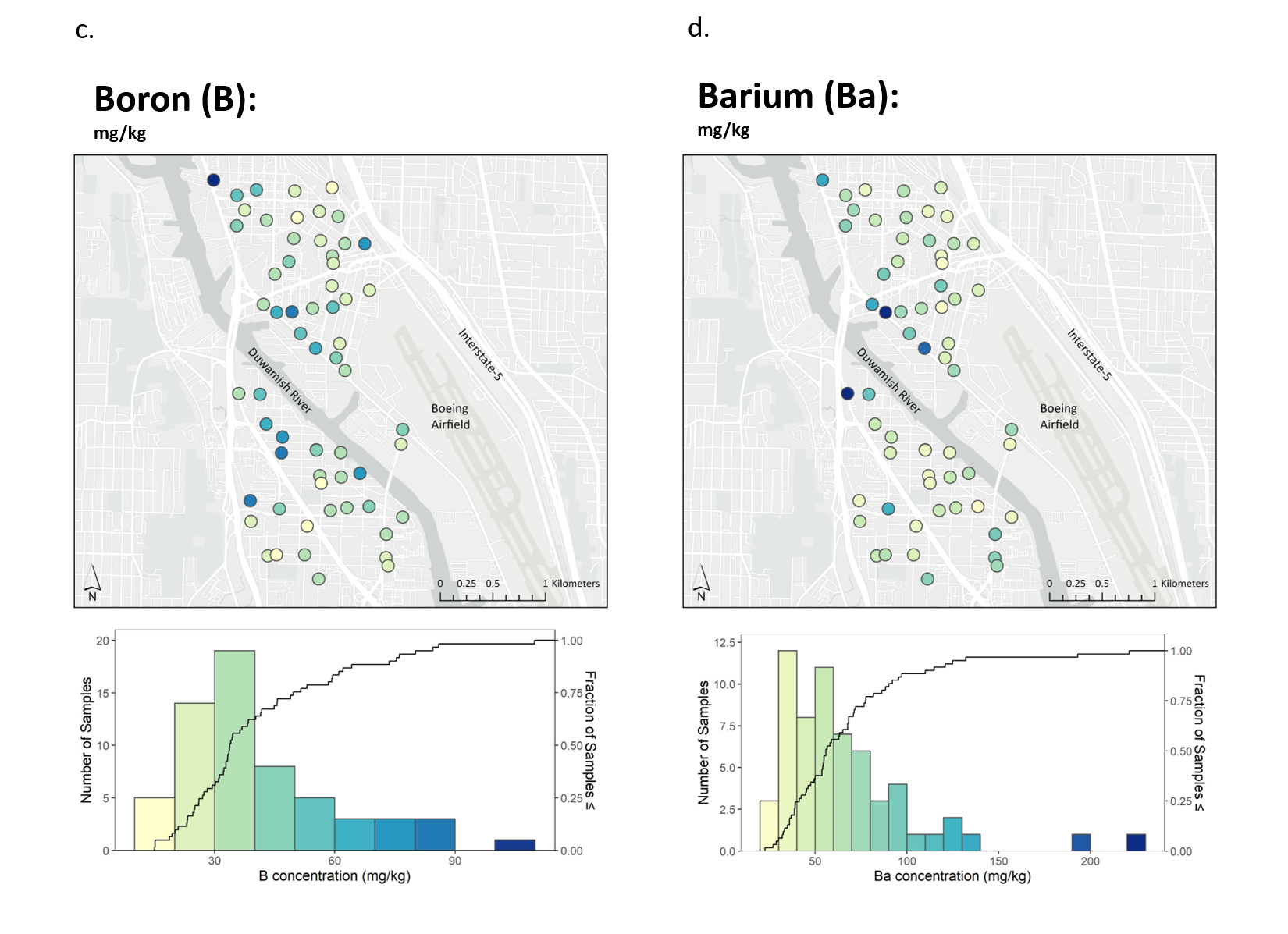

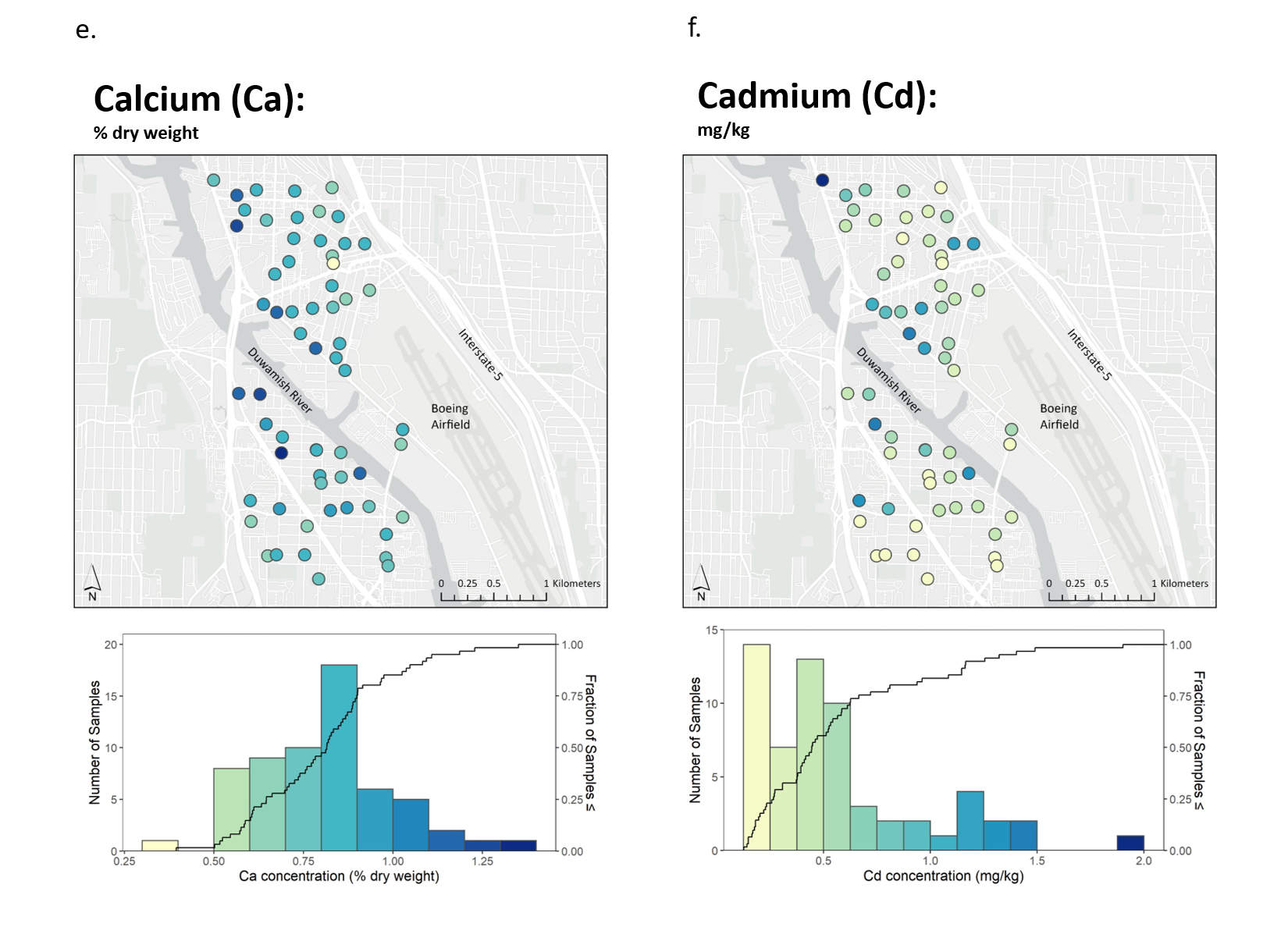

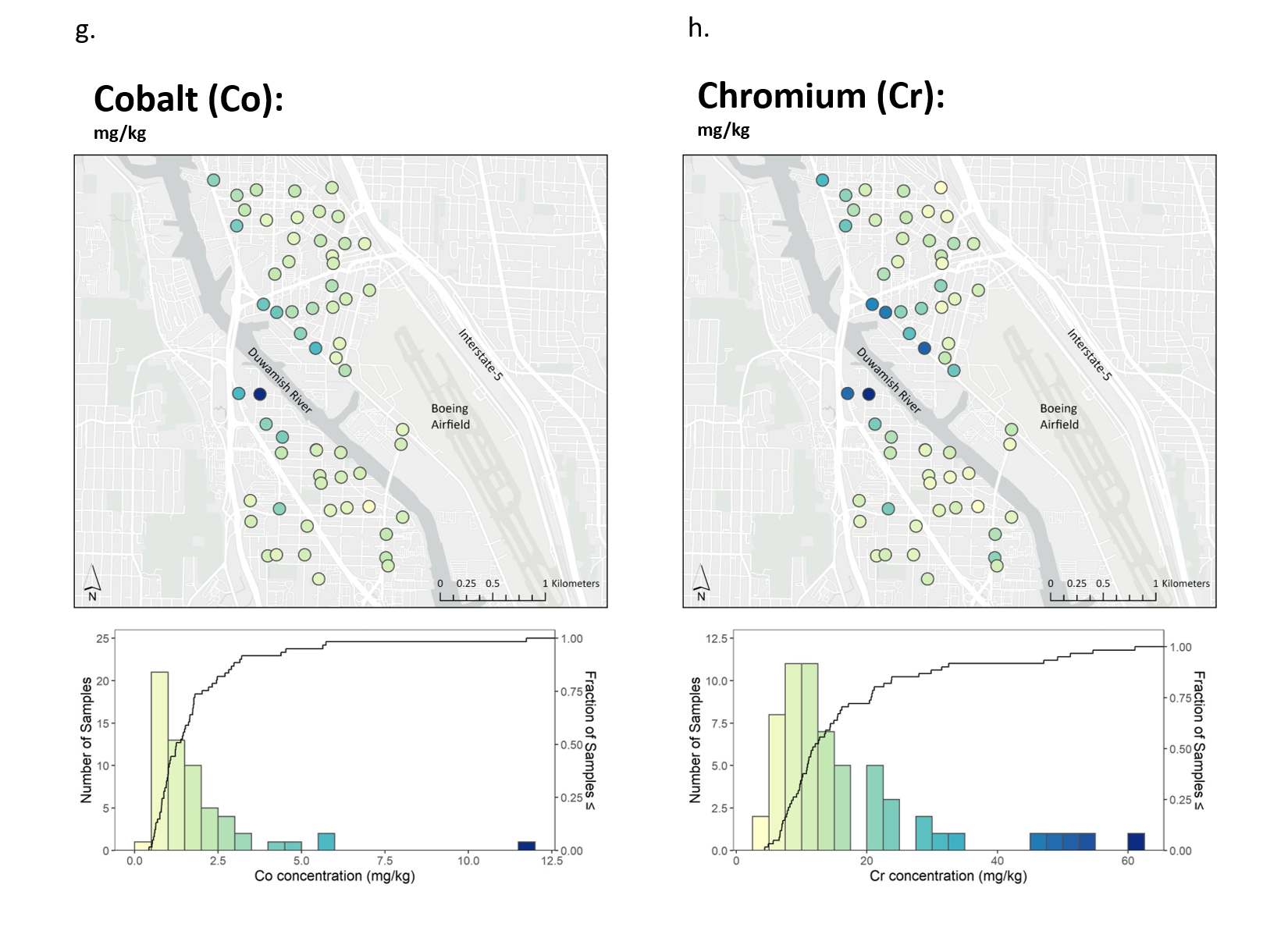

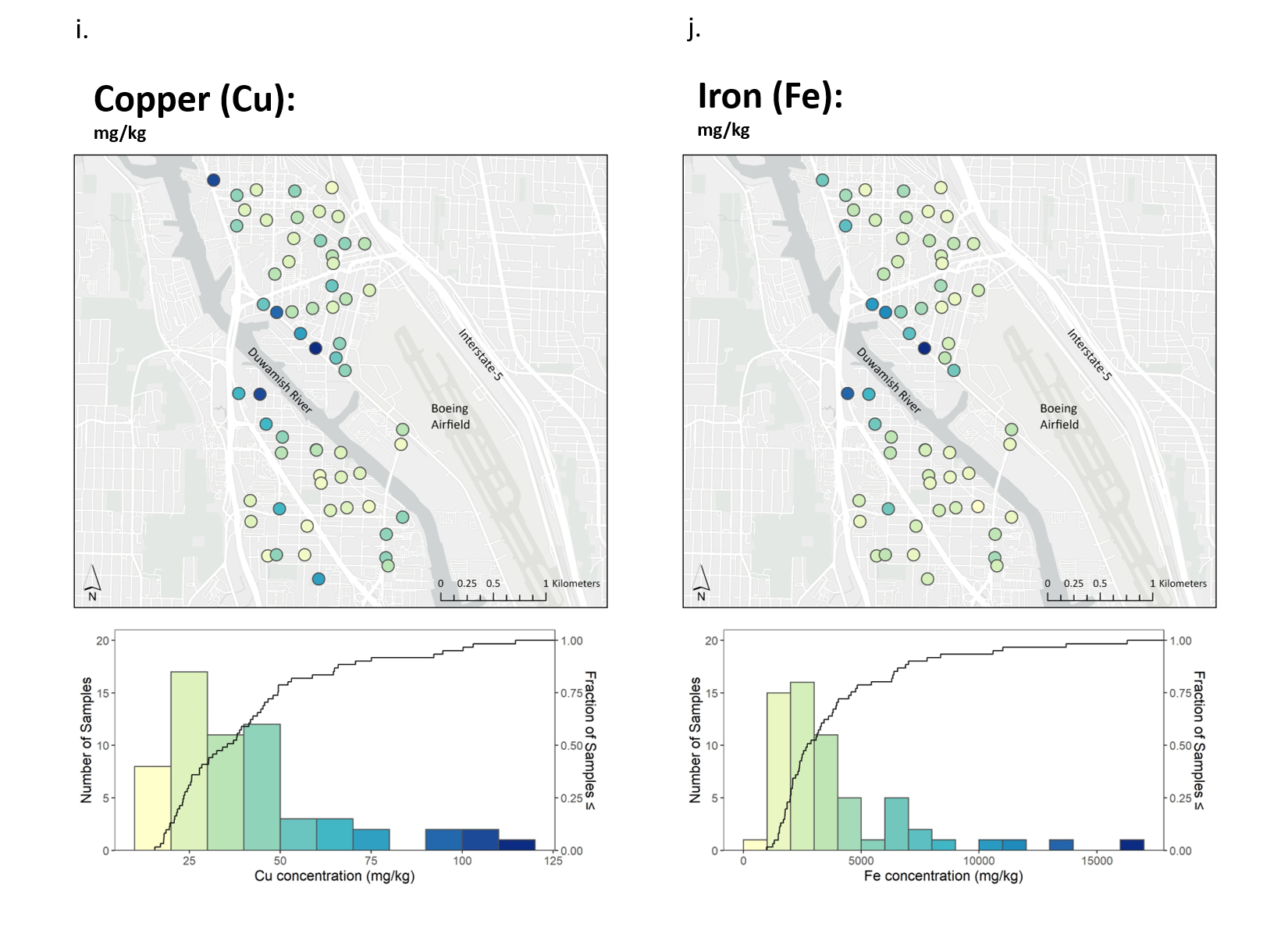

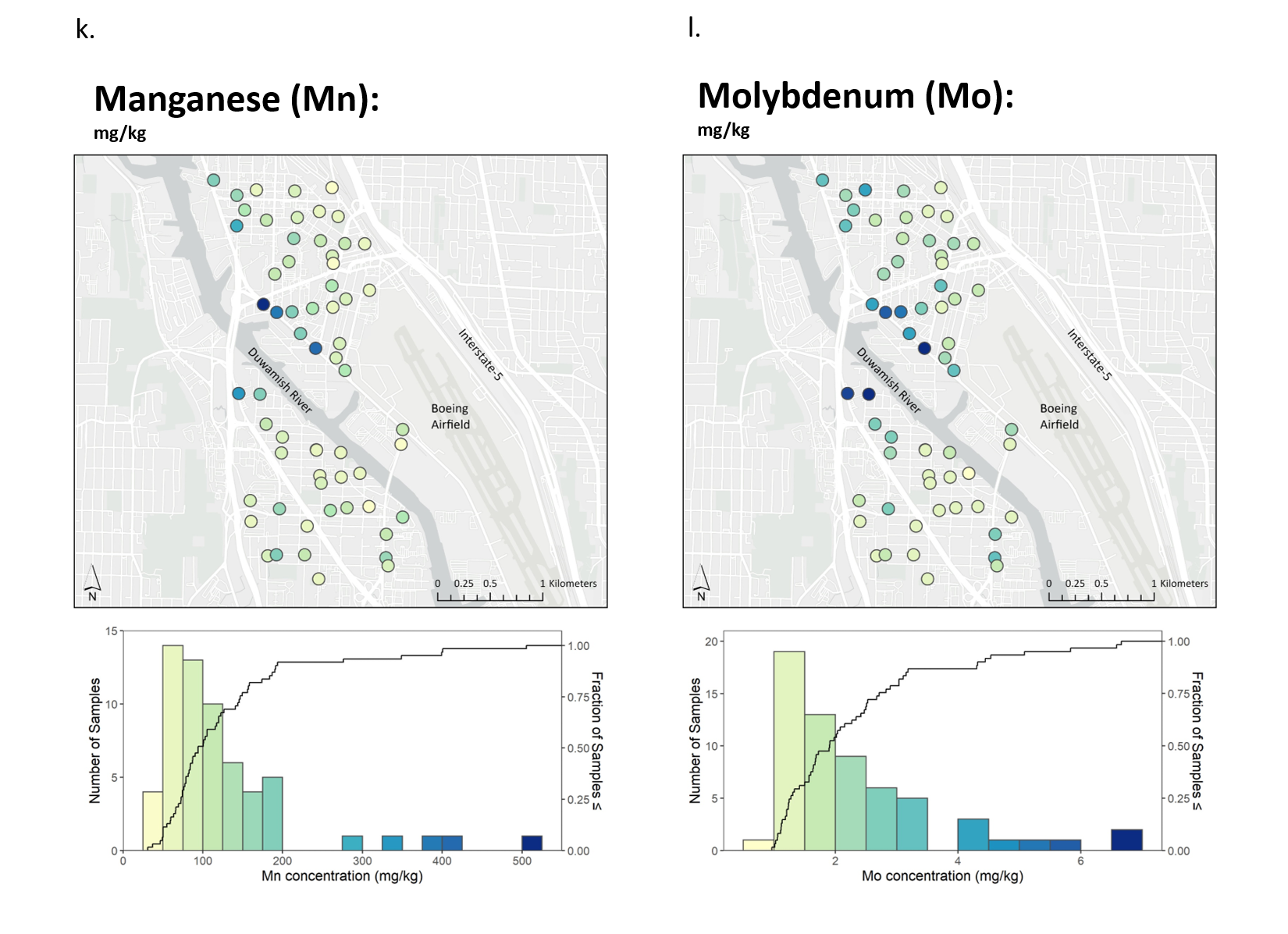

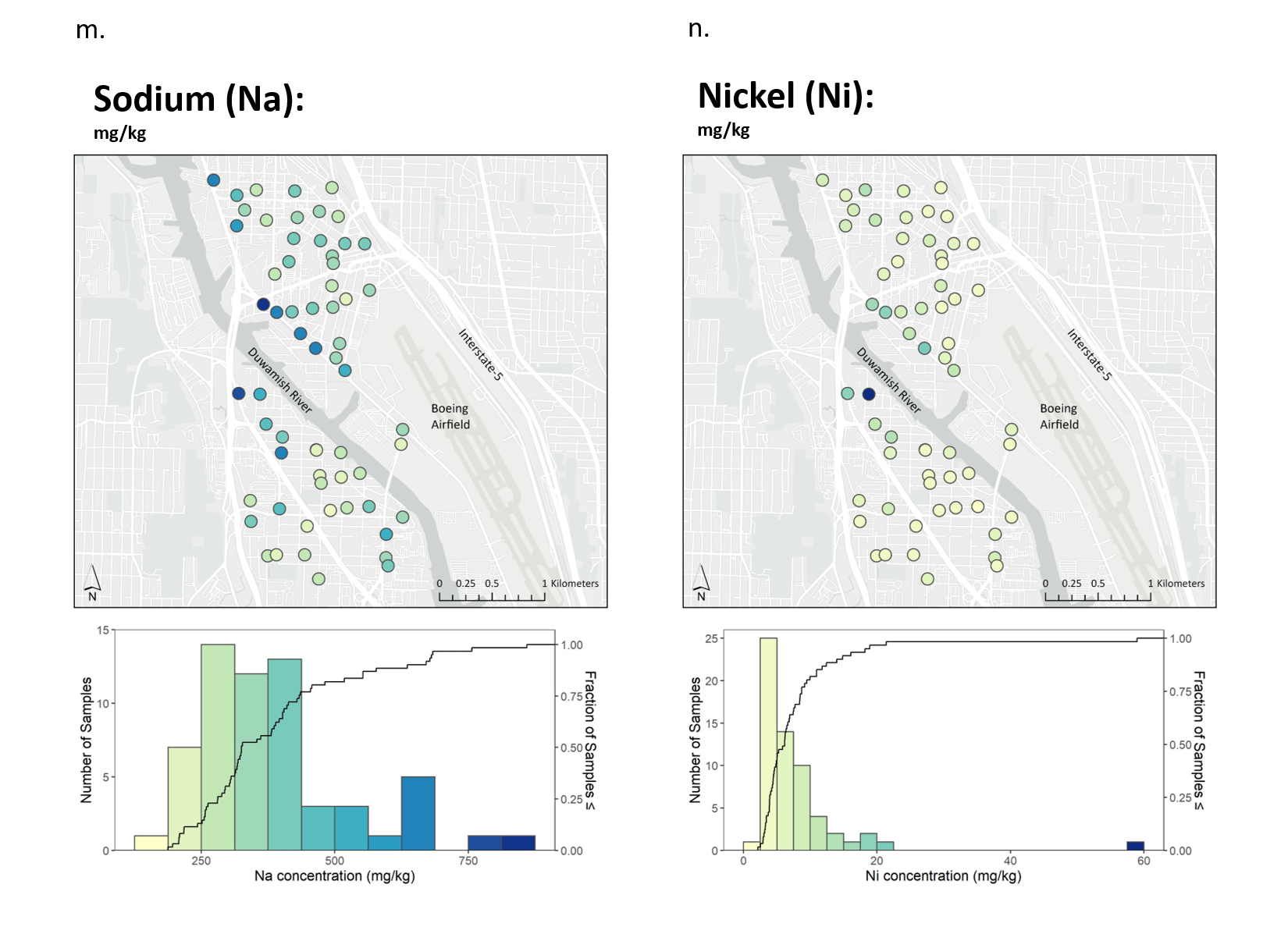

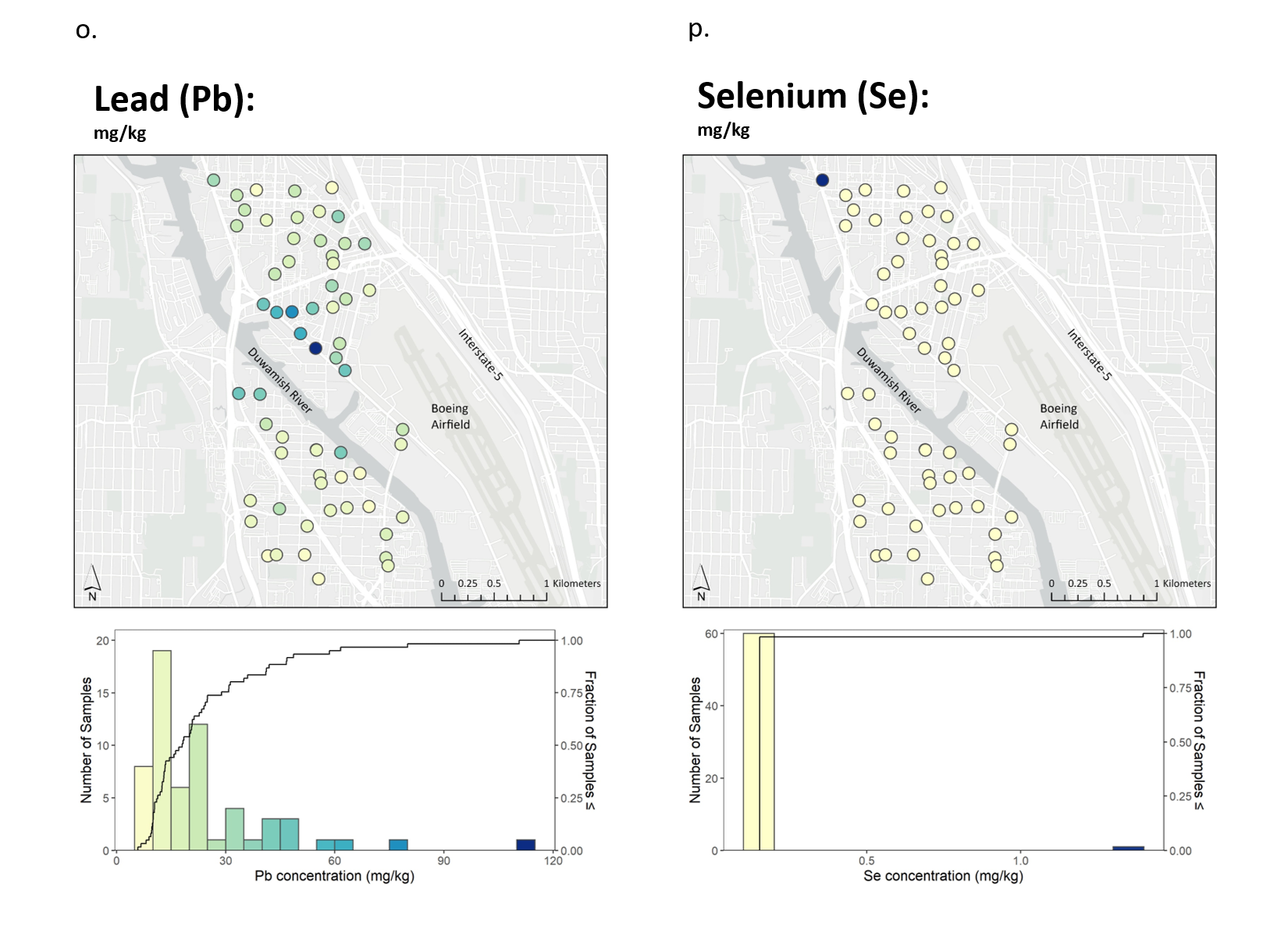

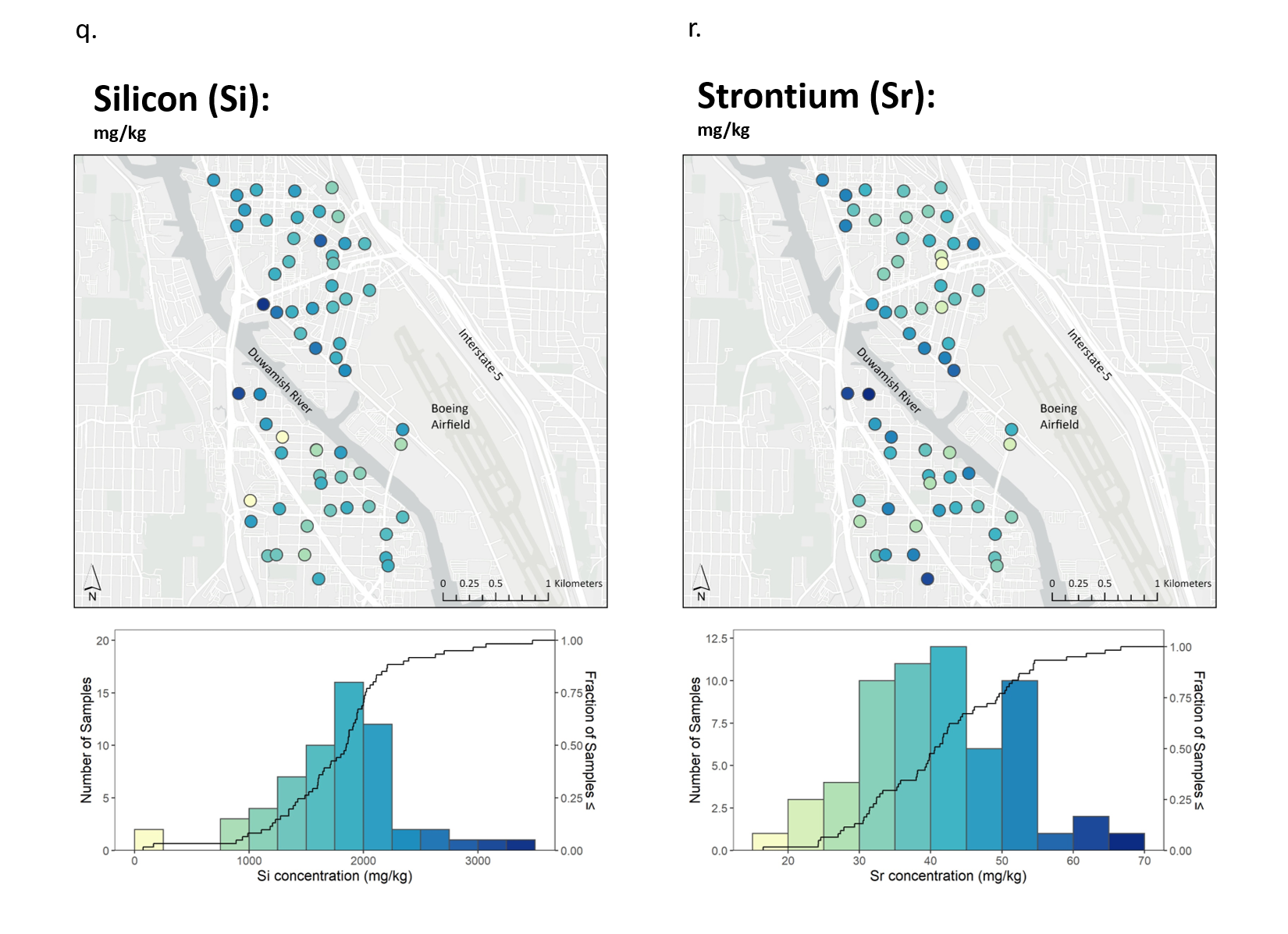

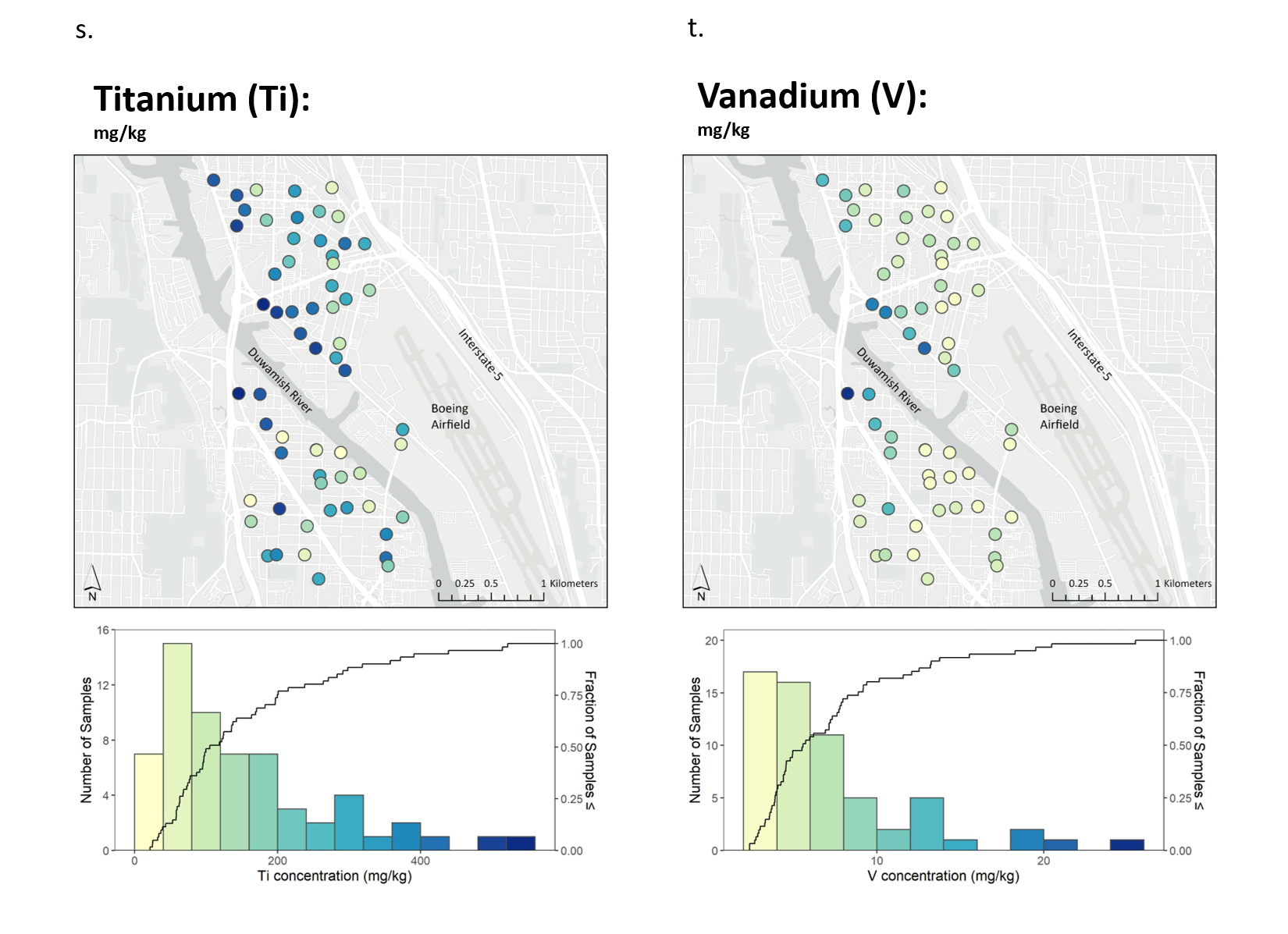

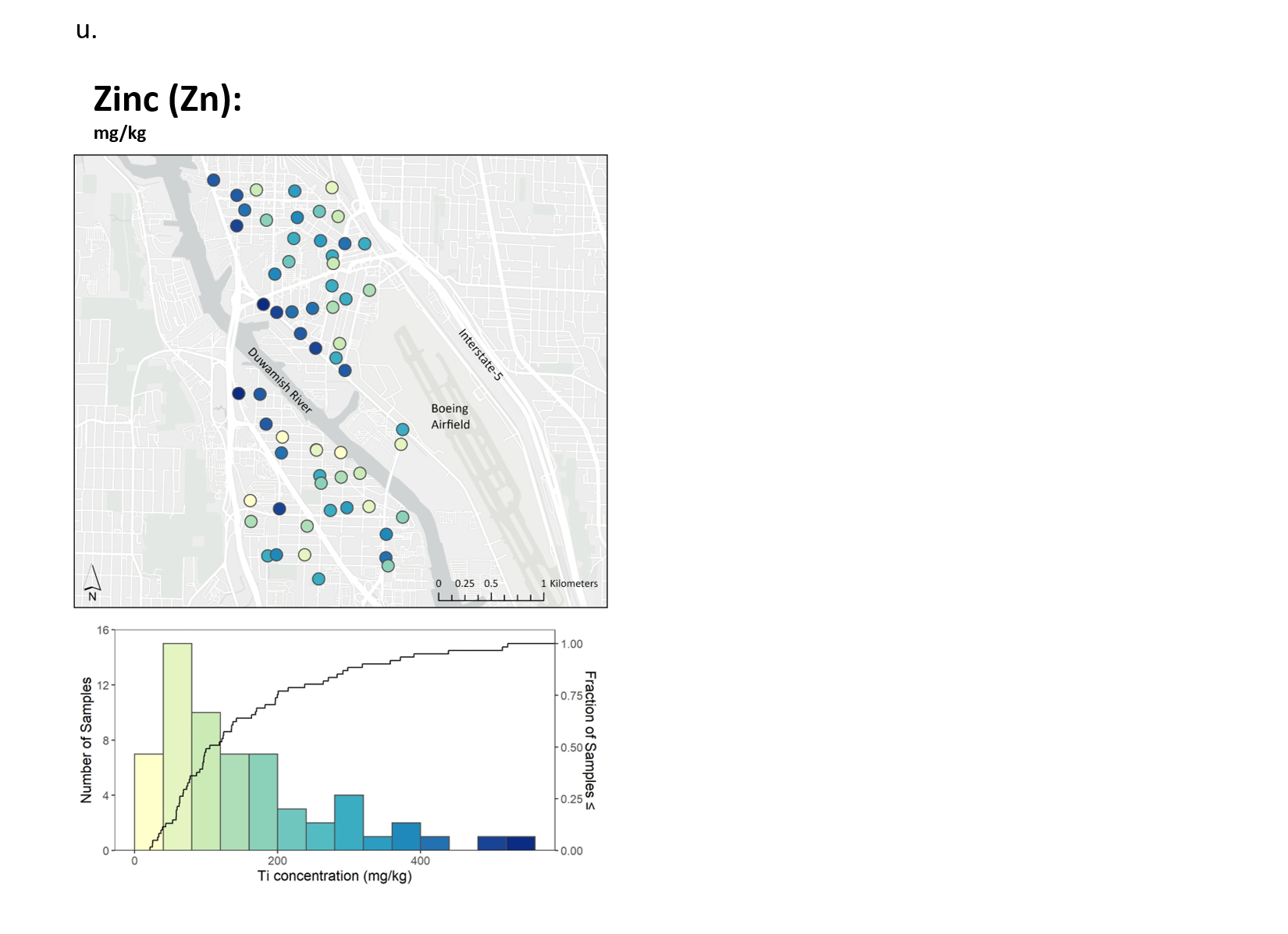
 Figure B.2: Dot maps and histograms showing concentrations of 21 chemical elements measured in moss, summer 2019. Black lines on the histograms are cumulative distribution curves.

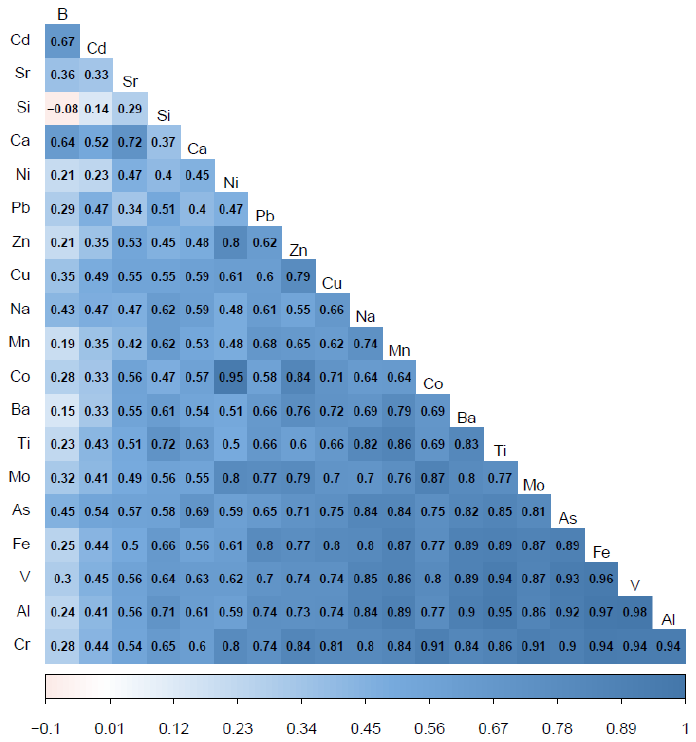

Figure B.3: Matrix of Pearson correlations between all elements measured in moss except for macronutrients of limited importance and selenium, for which concentrations in most samples were below detection limits (see Table 1).

1. As dictated by established standards, detection limits for plant-essential macro and secondary nutrients are reported in mg/kg and their concentrations in percent of dry weight
2. As dictated by established standards, detection limits for plant-essential macro and secondary nutrients are reported in mg/kg and their concentrations in percent of dry weight
